# A key role for phosphorylation of PsbH in the biogenesis and repair of photosystem II in Chlamydomonas

**DOI:** 10.1101/754721

**Authors:** Alexis Riché, Linnka Lefebvre-Legendre, Michel Goldschmidt-Clermont

## Abstract

Phosphorylation of the core subunits of photosystem II (PSII) is largely governed by a protein kinase and an antagonistic protein phosphatase. In plants the respective mutants show alterations in the architecture of thylakoid membranes and in the repair of PSII after photo-inhibition. However the protein kinase targets several subunits of PSII, as well as other proteins. To specifically investigate the role of phosphorylation of the different PSII subunits, we used site-directed mutagenesis and chloroplast transformation in *Chlamydomonas reinhardtii*. Major, evolutionarily-conserved sites of phosphorylation in three components of PSII (CP43, D2 and PsbH) were mutated to replace the corresponding serine or threonine residues with alanine. The alanine substitution mutant of D2 had no apparent phenotype, while the mutant of CP43 presented a minor delay in recovery from photo-inhibition. Alanine substitutions of the phosphorylation sites in PsbH had significant effects on the accumulation of PSII or on its recovery from photo-inhibition. When mutations in two of the target subunits were combined through a second cycle of chloroplast transformation, the strongest phenotype was observed in the mutant lacking phosphorylation of both PsbH and CP43, which showed delayed recovery from photo-inhibition. Surprisingly this phenotype was reversed in the mutant defective for phosphorylation of all three subunits. Our analysis indicates a prominent role for the N-terminus of PsbH in the stable accumulation of PSII and of PsbH phosphorylation in its repair cycle.

**SIGNIFICANCE STATEMENT:** To specifically investigate the role of PSII phosphorylation, alanine-substitution mutants of the major phospho-sites in the subunits of PSII were generated individually or in combinations using chloroplast transformation. PSII assembly was defective in some of the PsbH mutants. PSII repair after photo-inhibition was delayed most strongly in the mutant lacking phosphorylation of both PsbC (CP43) and PsbH.

## INTRODUCTION

Protein phosphorylation is a post-translational protein modification that plays important roles in the regulation of photosynthesis (Allen 1992). Some components of the photosynthetic machinery are prominent because of their abundance and their high degree of phosphorylation (Bennett 1980, Bennett 1984). These include components of Light Harvesting Complex II (LHCII) as well as subunits of the core of Photosystem II (PSII) such as PsbA (D1), PsbC (CP43), PsbD (D2) and PsbH, (Vener *et al.* 2001). The degree of phosphorylation of these proteins changes in response to environmental clues, notably the intensity and spectral quality of light, or in response to the metabolic state of the cell (Aro and Ohad 2003, Grieco *et al.* 2016, Wollman 2001). These changes reflect the role of protein phosphorylation in the regulation of light harvesting, of photosynthetic electron flow, and of the architecture of the thylakoid membrane network (Rochaix 2014). In plants the main enzymes that control thylakoid protein phosphorylation are the protein kinases STN7 and STN8 (STATE TRANSITION 7 and STATE TRANSITION 8), and the protein phosphatases PPH1/TAP38 (PROTEIN PHOSPHATASE 1 / THYLAKOID ASSOCIATED PHOSPHATASE 38) and PBCP (PHOTOSYSTEM II CORE PHOSPHATASE) (Bellafiore *et al.* 2005, Bonardi *et al.* 2005, Depege *et al.* 2003, Pribil *et al.* 2010, Samol *et al.* 2012, Vainonen *et al.* 2005). Although there is some degree of overlap in their substrate specificities, STN7 and PPH1/TAP38 mainly determine the phosphorylation of LHCII, while STN8 and PBCP primarily regulate the phosphorylation of the PSII core in Arabidopsis. Nevertheless it is only in the double mutant *stn7stn8* that phosphorylation of the major thylakoid proteins is minimal, while residual phosphorylation can be observed in the single mutants of the two kinases (Bonardi, et al. 2005).

The photosynthetic electron transfer chain is embedded in the thylakoid membranes of the chloroplast (Eberhard *et al.* 2008). In the linear mode of electron flow, PSII and Photosystem I (PSI) work in series to oxidize water and reduce ferredoxin (Fd). The reducing equivalents from Fd can then drive the conversion of NADP^+^ to NADPH, or be used in other metabolic pathways. The proton gradient that is concomitantly accumulated energizes ATP synthase, to produce ATP from ADP and Pi. In plants and green algae, light capture by PSI and PSII is enhanced by the respective light harvesting complexes LHCI and LHCII. In *Chlamydomonas reinhardtii*, dimers of PSII form supercomplexes by associating with monomeric “minor” LHCII subunits (LHCB4 and LHCB5) and LHCII trimers composed of a variety of isoforms (LHCBM1 to LHCBM9)(Drop *et al.* 2014). In plants such as Arabidopsis, the PSII supercomplex contains three types of monomeric LHCII, and trimers of three isoforms (Lhcb1, Lhcb2 and Lhcb3)(Galka *et al.* 2012, Su *et al.* 2017). In the alga, PSI forms supercomplexes by binding ten LHCI antenna subunits (Kubota-Kawai *et al.* 2019, Ozawa *et al.* 2018), while in plants the PSI-LHCI supercomplex contains only four LHCI subunits (Mazor *et al.* 2015).

In both plants and green algae such as *C. reinhardtii* (hereafter Chlamydomonas) a mobile part of LHCII can serve as an antenna for either PSII or PSI. Under low light its dynamic distribution is reflected in state transitions, a mechanism that regulates the allocation of LHCII to the two photosystems and maintains the redox poise of the electron transfer chain (Allen 1992, Rochaix 2014, Wollman 2001). Under high light, the energy-dependent component of non-photochemical quenching (qE) takes a major role in the regulation of photosynthetic electron flow by favoring the dissipation of excess energy as heat (Allorent *et al.* 2013, Ebenhoh *et al.* 2014, Niyogi 1999).

The phosphorylation of LHCII is a key determinant of state transitions. The protein kinases STT7 (in Chlamydomonas where it was first identified) or STN7 (in Arabidopsis) catalyze the phosphorylation of the mobile part of LHCII and induce its binding to PSI (Bellafiore, et al. 2005, Depege, et al. 2003). In conditions promoting state 2, when the electron transfer chain is over-reduced, the protein kinase is activated by the docking of reduced plastoquinol at the Q_o_ site of cytochrome b_6_f complex (Dumas *et al.* 2017, Vener *et al.* 1997, Zito *et al.* 1999). In plants, the Lhcb1 and Lhcb2 subunits of trimeric LHCII are the major substrates of STN7, and are phosphorylated on Thr residues close to their N-terminus (reviewed by (Pesaresi *et al.* 2011)). Phosphorylation of the Lhcb2 isoform is a central determinant for the binding of LHCII trimers to PSI and the formation of stable PSI-LHCI-LHCII super-complexes (Crepin and Caffarri 2015, Longoni *et al.* 2015, Pan *et al.* 2018). Recent evidence suggests that multiple LHCII trimers may bind PSI-LHCI at more than one site (reviewed by (Johnson and Wientjes 2019)). Under conditions where the PQ pool is oxidized that favor state 1, the protein phosphatase PPH1/TAP38 dephosphorylates LCHII and promotes the binding of the mobile trimers of LHCII to PSII (Pribil, et al. 2010, Shapiguzov *et al.* 2010). Thus phosphorylation of LHCII regulated by the antagonistic activities of STN7 and PPH1/TAP38 maintains a homeostatic redox poise of PQ and PQH2, by adjusting the antenna cross sections of PSII and PSI (Goldschmidt-Clermont and Bassi 2015).

While the role of LHCII phosphorylation is relatively well understood, less is known on the physiological function of PSII phosphorylation. In Arabidopsis, phosphorylation of the core subunits of PSII is mainly governed by the protein kinase STN8 and the phosphatase PBCP (Nath *et al.* 2013, Samol, et al. 2012, Vainonen, et al. 2005). However it is important to bear in mind that STN8 is involved in the phosphorylation of proteins other than the core subunits of PSII (Armbruster *et al.* 2013, Betterle *et al.* 2017, Reiland *et al.* 2011, Schonberg *et al.* 2017, Vainonen *et al.* 2008). The mutants *stn8* and *pbcp* show an altered organization of thylakoid membranes. In *stn8*, and particularly in the double mutant *stn7stn8*, the diameter of the thylakoid grana is larger compared to the wild type (Fristedt *et al.* 2009, Iwai *et al.* 2018, Nath, et al. 2013). In contrast in the *pbcp* phosphatase mutant, the diameter of the grana is similar to the wild type, but the number of membrane layers in grana stacks is reduced (Samol, et al. 2012). State transitions are not affected in the *stn8* or *pbcp* mutants (Bonardi, et al. 2005, Samol, et al. 2012).

Light causes damage to PSII, and in conditions where the rate of damage exceeds the rate of repair, photo-inhibition becomes apparent. The primary site of PSII damage is the D1 subunit (PsbA), which can be selectively degraded and replaced by a newly synthetized subunit (Schuster *et al.* 1988). Because PSII localizes to the stacked grana regions, and repair takes place in the stroma-exposed lamella, the repair cycle is thought to involve the dismantling of PSII super-complexes and the migration of damaged PSII from grana to stromal lamellae (reviewed by (Aro *et al.* 2005, Jarvi *et al.* 2015, Komenda *et al.* 2012, Theis and Schroda 2016)). The *stn8* and *pbcp* mutants show enhanced sensitivity to photo-inhibition and delayed repair of damaged PSII (Nath, et al. 2013, Puthiyaveetil *et al.* 2014b, Tikkanen *et al.* 2008). To explain these phenotypes of *stn8* and *pbcp* there are two main types of hypotheses, which are not mutually exclusive. The first is that changes in thylakoid membrane stacking due to altered phosphorylation of the targets of STN8 and PBCP could influence the efficiency of repair indirectly, by modulating the access of damaged PSII to the repair machinery (Fristedt, et al. 2009, Puthiyaveetil *et al.* 2014a, Puthiyaveetil *et al.* 2017). Phosphorylation could also affect the lateral mobility of thylakoid proteins within the membrane network (Goral *et al.* 2010, Herbstova *et al.* 2012). The alternative type of hypotheses propose that the sensitivity to photoinhibition of *stn8* and *pbcp* could be due to a direct requirement for PSII protein phosphorylation and subsequent de-phosphorylation in the repair cycle of the D1 subunit (Baena-Gonzalez *et al.* 1999). Biochemical analysis of photoinhibition in intact plant chloroplasts or isolated thylakoids showed that phosphorylated D1 and D2 are degraded more slowly than the non-phosphorylated forms (Ebbert and Godde 1996, Koivuniemi *et al.* 1995). Degradation of damaged D1 involves chloroplast proteases of the FtsH and Deg families (Bailey *et al.* 2002, Kapri-Pardes *et al.* 2007, Kato and Sakamoto 2009, Lindahl *et al.* 2000, Malnoe *et al.* 2014). In the *var2* mutant of Arabidopsis, which is deficient for the protease FtsH2 and hyper-sensitive to photoinhibition, phosphorylation of D1 is increased and its degradation is delayed (Kato and Sakamoto 2014). However these phenotypes are suppressed in the double mutant *var2 stn8*, implicating a role for protein phosphorylation in the control of D1 degradation.

In Chlamydomonas, the major phosphorylated subunits of PSII were originally identified by labelling *in vivo* with ^32^P-phosphate, or by immuno-blotting with antisera against phospho-threonine (de Vitry *et al.* 1991, Delepelaire 1984, Delepelaire and Wollman 1985). Subsequently numerous phosphorylation sites in the subunits of PSII were identified by mass spectrometry (reviewed by (Grieco, et al. 2016)). After labelling with radioactive ^32^P-phosphate, the major phosphoproteins of PSII were found to be PsbC (CP43), PsbD (D2) and PsbH (de Vitry, et al. 1991, Delepelaire 1984). D1 phosphorylation was not detected by ^32^P-phosphate labelling, although it was observed by mass spectrometry (Turkina *et al.* 2006, Wang *et al.* 2014). In contrast D1 is a major phosphoprotein in seed plants (Chen *et al.* 2018, Pursiheimo *et al.* 1998), but not in the moss *Physcomitrella patens* (Gerotto *et al.* 2019). In Chlamydomonas the extent of phosphorylation of most of these proteins is not known, with the exception of D2. Its phosphorylated form, which migrates slightly more slowly during SDS-PAGE, is more abundant than the non-phosphorylated form under certain conditions, indicating that the protein can undergo a large degree of phosphorylation (Delepelaire 1984).

Many studies on the role of PSII phosphorylation are based on the phenotypes of *stn8* and *pbcp*, with the tacit assumption that the mutant phenotypes are due to the altered phosphorylation of the core subunits of the complex. However phosphorylation of different subunits of PSII could have distinct functions. Furthermore phospho-proteomic studies have shown that there exist other targets of STN8 (Reiland, et al. 2011, Schonberg, et al. 2017). Examples of putatively relevant phosphorylation targets include the members of the CURT1 (CURVATURE THYLAKOID 1) protein family, which have strong effects on thylakoid architecture and are thought to facilitate membrane bending at the edges of grana stacks (Armbruster, et al. 2013). CAS (Calcium Sensor) phosphorylation might also have an effect, as the protein is involved in the regulation of qE in Chlamydomonas and is known to be a target of STN8 in Arabidopsis (Petroutsos *et al.* 2011, Vainonen, et al. 2008). Here we directly address the role of phosphorylation of the individual PSII core subunits in Chlamydomonas by using chloroplast transformation for site-directed mutagenesis of the major phosphorylation sites. We also examine the effect of cumulating mutations in two or three of the major phosphorylated subunits.

## RESULTS

### Site-directed mutagenesis of the major phosphorylation sites in PSII

For our investigations we selected five phosphorylation sites in three PSII subunits (Table 1), using criteria of strong phosphorylation and evolutionary conservation. For PsbC in Chlamydomonas, phosphorylation was most frequently observed at Thr3 (Lemeille *et al.* 2010, Turkina, et al. 2006, Wang, et al. 2014), and furthermore phosphorylation at this site was conserved during evolution of higher plants (reviewed by (Grieco, et al. 2016)), hence it was selected as a target of our analysis (Table 1). Two amino-acid residues of PsbC are removed post-translationally from its N-terminus, so that in the mature protein Thr3 is the N-terminal residue, which can furthermore be acetylated. Likewise for D2 (PsbD) Thr2 was selected for our analysis, as its phosphorylation was repeatedly observed and was evolutionarily conserved in higher plants (Grieco, et al. 2016, Lemeille, et al. 2010, Turkina, et al. 2006, Wang, et al. 2014). After post-translational processing, Thr2 is the N-terminal residue and can be acetylated. Phosphorylation of PsbH, also known as the phosphoprotein of 9 or 10 kDa, is more complex. In Chlamydomonas three sites of phosphorylation have been observed at Thr3, Thr5 and Ser6. Only di-phosphorylated peptides were observed, at Thr3 and Thr5, or at Thr3 and Ser6 (Lemeille, et al. 2010, Wang, et al. 2014). Phosphorylation at two of these sites, Thr3 and Thr5, was conserved during evolution in higher plants, where mono-phosphorylation at Thr3 and di-phosphorylation at Thr3 and Thr5 were observed (reviewed by (Grieco, et al. 2016)). However the third site, Ser 6, is replaced by Val in higher plants (Table 1). In contrast to seed plants, in Chlamydomonas D1 is not strongly phosphorylated and was therefore not included in this study.

**Table 1.**
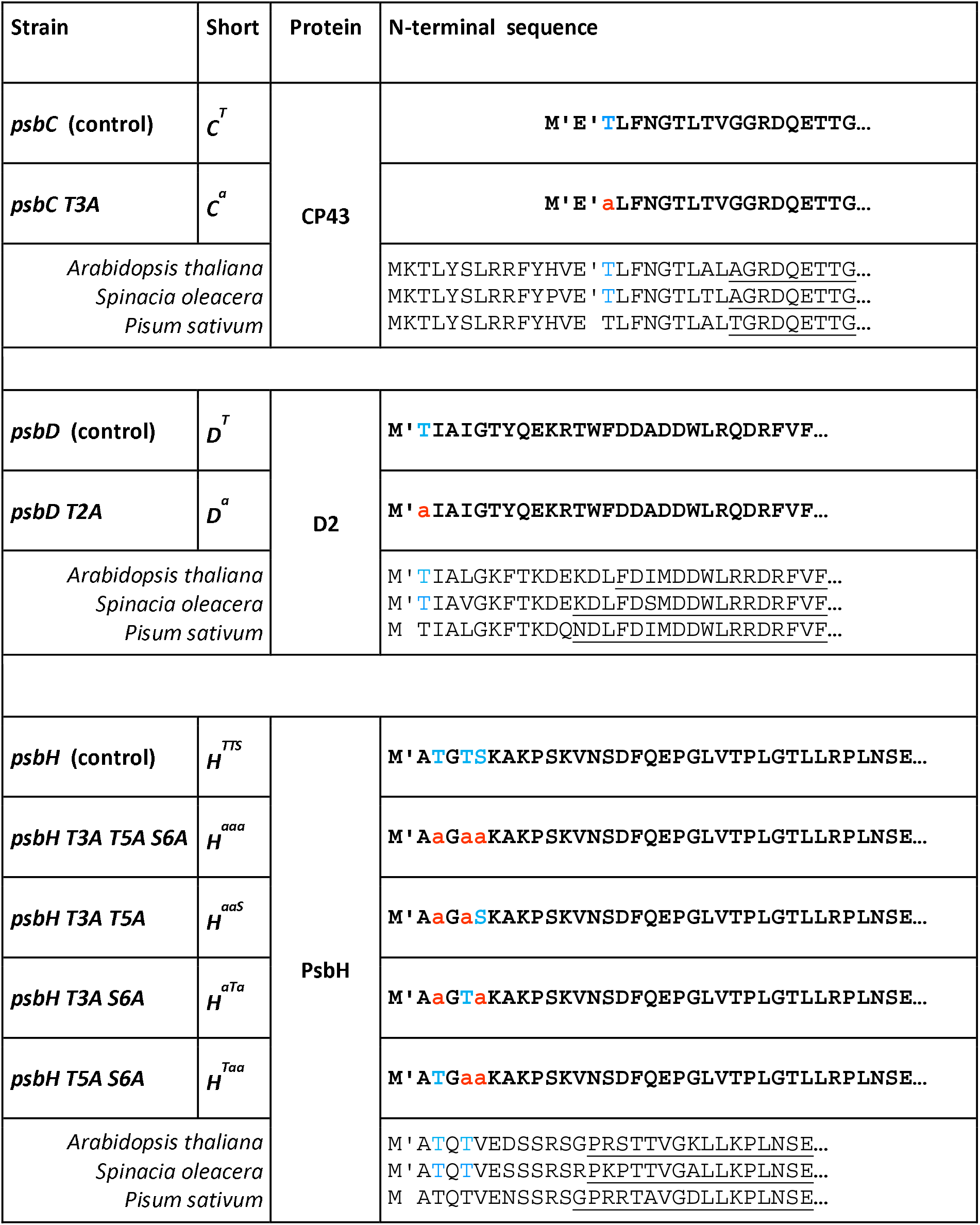
Alanine substitution mutants and the respective control strains used in this work. The table shows the correspondence between the full name and the shorthand name of the strain, the N-terminal sequence of the protein carrying the alanine substitutions (in red) compared to the wild-type sequence of the control, with the Thr or Ser phospho-sites shown in blue. The position of the N-terminus of the mature proteins after post-translational processing is shown with a tick (’). For CP43 (PsbC), two different N-terminal processing sites have been identified (Lemeille, et al. 2010, Wang, et al. 2014). The sequences of the N-terminal part of the orthologous proteins from *Arabidopsis thaliana*, *Spinacia oleacera* and *Pisum sativum* are shown for comparison. The phospho-sites identified by mass spectrometry are shown in blue (reviewed in (Grieco, et al. 2016)), but they have not to our knowledge been determined in *P. sativum*. The structured part of the N-terminal domain of the proteins, which is resolved in the PSII structures determined by cryo-electron-microscopy, is underlined (Su, et al. 2017, van Bezouwen *et al.* 2017, Wei *et al.* 2016). All strains additionally contain a footprint of the excisable *aadA* cassette at the respective locus (see Fig 1).

To investigate the role of protein phosphorylation at these major phosphorylation sites in PsbC, PsbD and PsbH, we generated site-directed mutants where the respective Thr or Ser residues were changed to Ala, a residue which cannot be phosphorylated (Table 1). Similar mutants of some of the phosphorylation sites have been previously described, namely Thr2 of PsbD (Andronis *et al.* 1998, Fleischmann and Rochaix 1999) and Thr3 of PsbH (O’Connor *et al.* 1998). A mutation of D2 was included in our analysis for completeness, and to allow the construction of lines combining several phospho-site mutations. For PsbH, we designed a mutant lacking all three phosphorylation sites, and three other mutants each retaining only one of the phosphorylation sites (Table 1). To facilitate the description of the mutants in this report, we have adopted shorthand acronyms where the mutated gene is designated with a single capital letter, such as *C* for *PsbC*. This is followed by a superscript indicating whether the encoded Thr or Ser residue is wild-type (uppercase T or S) or has been mutated to Ala (lowercase a). For example the mutant of *psbC* that encodes a protein where Thr2 is replaced by Ala (PsbC T2A) is abbreviated *C*^*a*^, while the corresponding control with the wild-type sequence is *C*^*T*^. For *psbH*, the three residues at the positions of Thr3, Thr5 and Ser6 are listed sequentially in the superscript. For example a mutant line with *psbH* mutated to encode two Ala residues in place of Thr3 and Ser6, but retain wild-type Thr5, is annotated *H*^*aTa*^.

For chloroplast transformation, three plasmid vectors were constructed carrying fragments of plastome DNA centered on the target sites of directed mutagenesis. To allow selection in Chlamydomonas, the *aadA* marker conferring spectinomycin resistance (Goldschmidt-Clermont 1991) was introduced in a non-coding region either upstream (*psbC*, *psbD*) or downstream (*psbH*) of the target gene (Fig 1). The *aadA* cassette was flanked by direct repeats (483 bp), so that the marker would be excisable through spontaneous homologous recombination (Fischer *et al.* 1996). The desired mutations were engineered into the transformation vectors (Supplemental Fig S1 and Table S1), which were then used for biolistic transformation of wild-type *C. reinhardtii*. The resulting transformants were repeatedly sub-cultured on selective spectinomycin-containing medium until genotyping showed that homoplasmy had been obtained (Supplemental Figs S2 and S3). The lines were then sub-cultured on medium lacking spectinomycin, to allow excision and loss of the *aadA* cassette, leaving behind only one copy (“footprint”) of the 483 bp repeat. The excision of the *aadA* cassette, verified by genotyping, allowed a second round of transformation to generate mutant combinations with alanine substitutions at two chloroplast loci (see below and Supplemental Fig S4). A third round of chloroplast transformation was finally used to obtain mutant lines with alanine substitution mutations at all three loci of interest. In all cases, control transformed lines carrying only 483 bp footprints at the corresponding loci, but with wild-type gene sequences, were obtained in parallel for direct comparisons in isogenic backgrounds.

**Figure 1.**
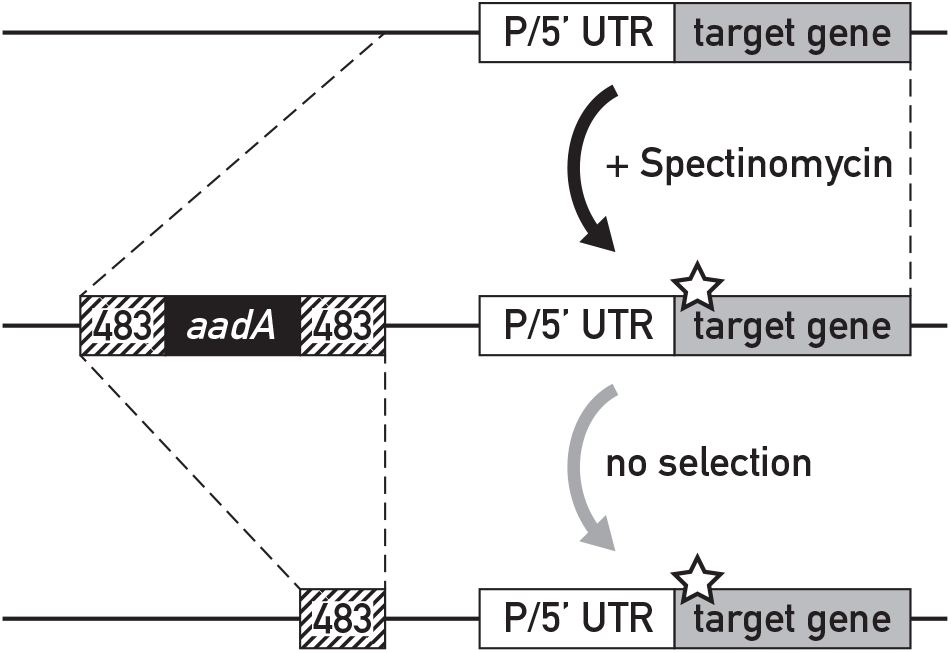
Transformation strategy. The transformation vector contained the excisable *aadA* selection cassette (flanked by 483 bp direct repeats) inserted in close proximity to the target. The desired site-directed mutation of the phosphorylation site is shown with a star. After biolistic chloroplast transformation and homologous recombination, transformants were selected on spectinomycin and subcultured repeatedly until homoplasmy was confirmed by genotyping. Loss of the *aadA* cassette by recombination between the 483 bp repeats was allowed by subculturing in the absence of spectinomycin until the strains were homoplasmic, as confirmed by genotyping.

### Mutations of the phosphorylation sites in PsbH affect PSII accumulation

The lines with alanine substitutions in PsbC, PsbD or PsbH and the corresponding controls were tested for their ability to grow on acetate-containing medium (TAP) in the dark, and on minimal medium (HSM) in the light (Suppl. Fig S5). All the strains were capable of heterotrophic growth in the dark as expected. The alanine-substitution mutants in PsbC (*C*^*a*^) and D2 (*D*^*a*^) grew photo-autotrophically on minimal medium similar to the wild type or the controls *C*^*T*^ and *D*^*T*^ (Supplemental Fig S5). This indicates that phosphorylation at Thr3 of CP43 or Thr2 of D2 is not essential for photosynthesis. Of the three PsbH mutants retaining only one phosphorylation site, *H*^*aaS*^ and *H*^*Taa*^ and the control *H*^*TTS*^ grew normally on minimal medium while in contrast *H*^*aTa*^ grew only very slowly. The triple mutant of PsbH (*H*^*aaa*^) failed to grow photo-autotrophically. These observations suggest that Thr3 and Ser6 have a redundant but essential function for photosynthesis that cannot be fully compensated by Thr5.

To monitor the activity of PSII, we used chlorophyll fluorescence spectroscopy and measured the maximum quantum yield of PSII (Fv/Fm) in the dark-adapted state and its quantum efficiency (ΦPSII) at different light intensities (Fig. 2A) (Maxwell and Johnson 2000). For this analysis the cells were grown on acetate-containing medium in the dark. All the control lines as well as *C*^*a*^, *D*^*a*^ and *H*^*aaS*^ had a maximum quantum yield similar to the wild type (Fv/Fm = 0.65 ~ 0.70). In contrast, *H*^*aaa*^ and *H*^*aTa*^ had very low values, consistent with their failure to grow photoautotrophically (Suppl. Fig. S5). Unexpectedly, *H*^*Taa*^ was strongly affected, with Fv/Fm = 0.28, although it was capable of normal growth on minimal medium in the light. However when *H*^*Taa*^ was cultivated in the light (60 μE m^−2^ s^−1^), it had an Fv/Fm value of 0.70 (Fig S6). It thus appears that after cultivation in the light *H*^*Taa*^ is photosynthetic with active PSII, but that after cultivation in the dark its PSII complex is either partly inactive, or less abundant relative to LHCII. Data presented in the next paragraph argue in favor of the latter alternative. The quantum efficiency of PSII was measured at three levels of light intensity in those lines that retained significant or normal values of Fv/Fm after growth in the dark (Fig. 2B). The mutants *C*^*a*^, *D*^*a*^ and *H*^*aaS*^ had ΦPSII values similar to the controls, while *H*^*Taa*^ had lower values that paralleled its defect in Fv/Fm.

**Figure 2.**
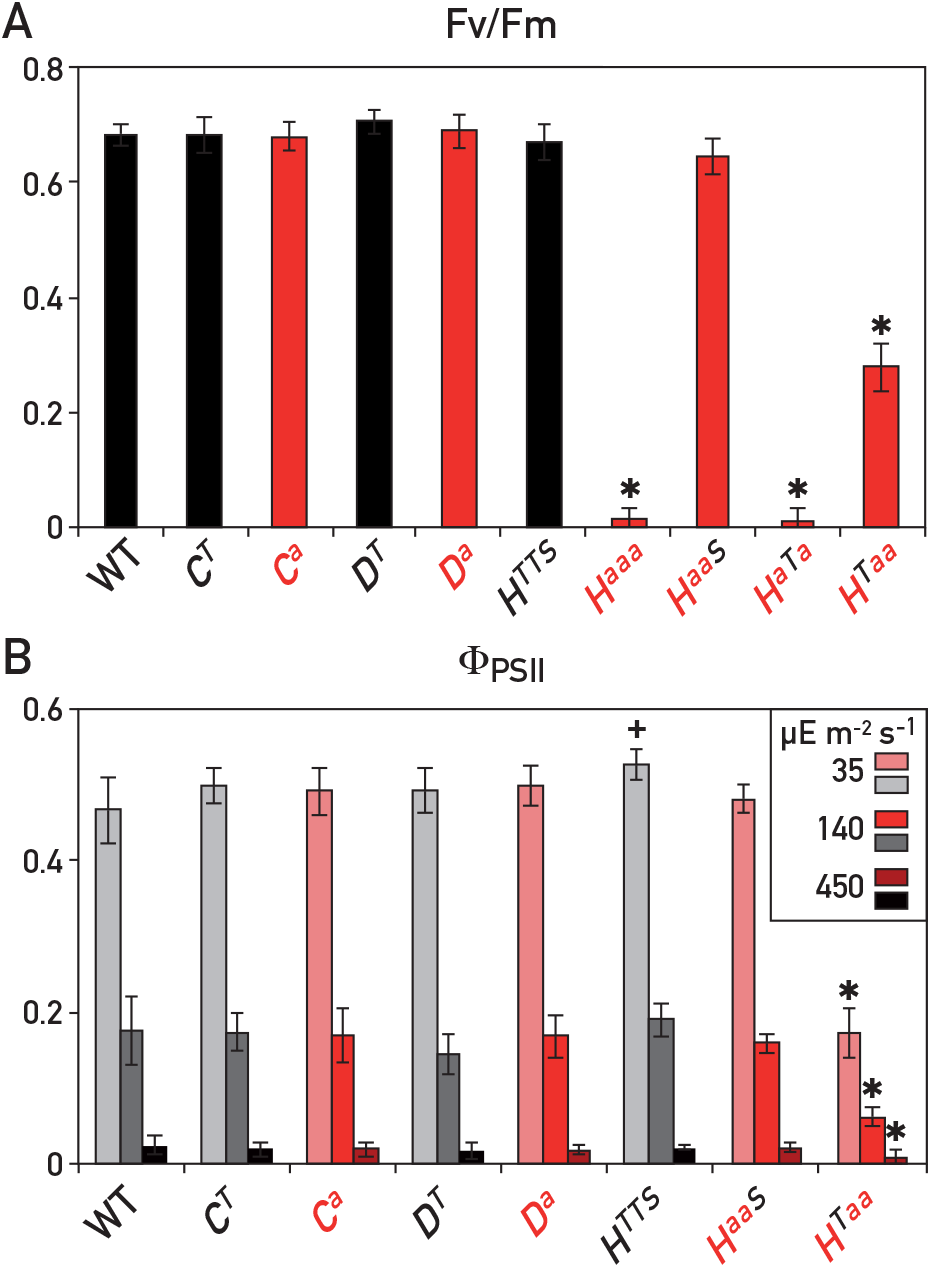
Chlorophyll fluorescence spectroscopy. A. Maximum quantum yield of photosystem II (F_v_/F_m_ = (F_m_-F_o)_/F_m_). Cells were grown in TAP medium in the dark. B. Quantum yield of photosystem II (ϕ_PSII_ = (F_m_’-F_t_)/F_m_’) at different light intensities (32, 140 or 450 μE m^−2^ s^−1^). Cells were grown in the dark in acetate medium (TAP), transferred to minimal medium (HSM) and incubated for 1h in low light (6 μE.m^−2^.s^−1^) before aliquots of the cells were exposed to the specified light intensity for 5 minutes to allow ϕ_PSII_ to stabilize. Each value is the average of three measurements on two independent lines of identical genotype (+/− SD). Significant differences in Student’s t-test between the control strains and the wild type, or between the mutants and their respective controls, are indicated (+: p<0.05; *: p<0.01).

### Light-dependent assembly of PSII in the *H*^*Taa*^ mutant of PsbH

The accumulation of PSII and other thylakoid protein complexes was monitored in total protein extracts by SDS-PAGE and immunoblotting (Fig 3). A mutant deficient for the D1 subunit (*Fud7*, (Bennoun *et al.* 1986)) was also included as the absence of this core subunit of PSII also destabilizes the entire complex. For this analysis the mutant lines and controls were grown in the dark. The immunoblots were decorated with antisera against the mutated subunit (CP43, D2 or PsbH respectively) as well as D1 (representative of PSII), PsaA (PSI), Lhcbm5 (LHCII) and AtpB (ATP synthase). The control strains and the *C*^*a*^, *D*^*a*^ and *H*^*aaS*^ mutants had normal amounts of the PSII subunits, and also of the components of the other complexes. Instead *H*^*aaa*^ and *H*^*aTa*^ had undetectable amounts of PsbH and very low levels of D1. Nevertheless they accumulated the other photosynthetic complexes to normal levels. Thus in these two genotypes PSII is specifically absent. Dark-grown *H*^*Taa*^ had only low levels of PsbH and D1 (approx. 30% of the wild type), while light-grown *H*^*Taa*^ displayed levels of these subunits similar to the wild type or the *H*^*TTS*^ control. These observations are consistent with the respective values of Fv/Fm (Fig 2 and Supplemental Fig S6) after cultivation in the dark (0.28) or in the light (0.70). They imply that PSII accumulation is affected in dark-grown *H*^*Taa*^ but that this defect is alleviated in the light.

**Figure 3.**
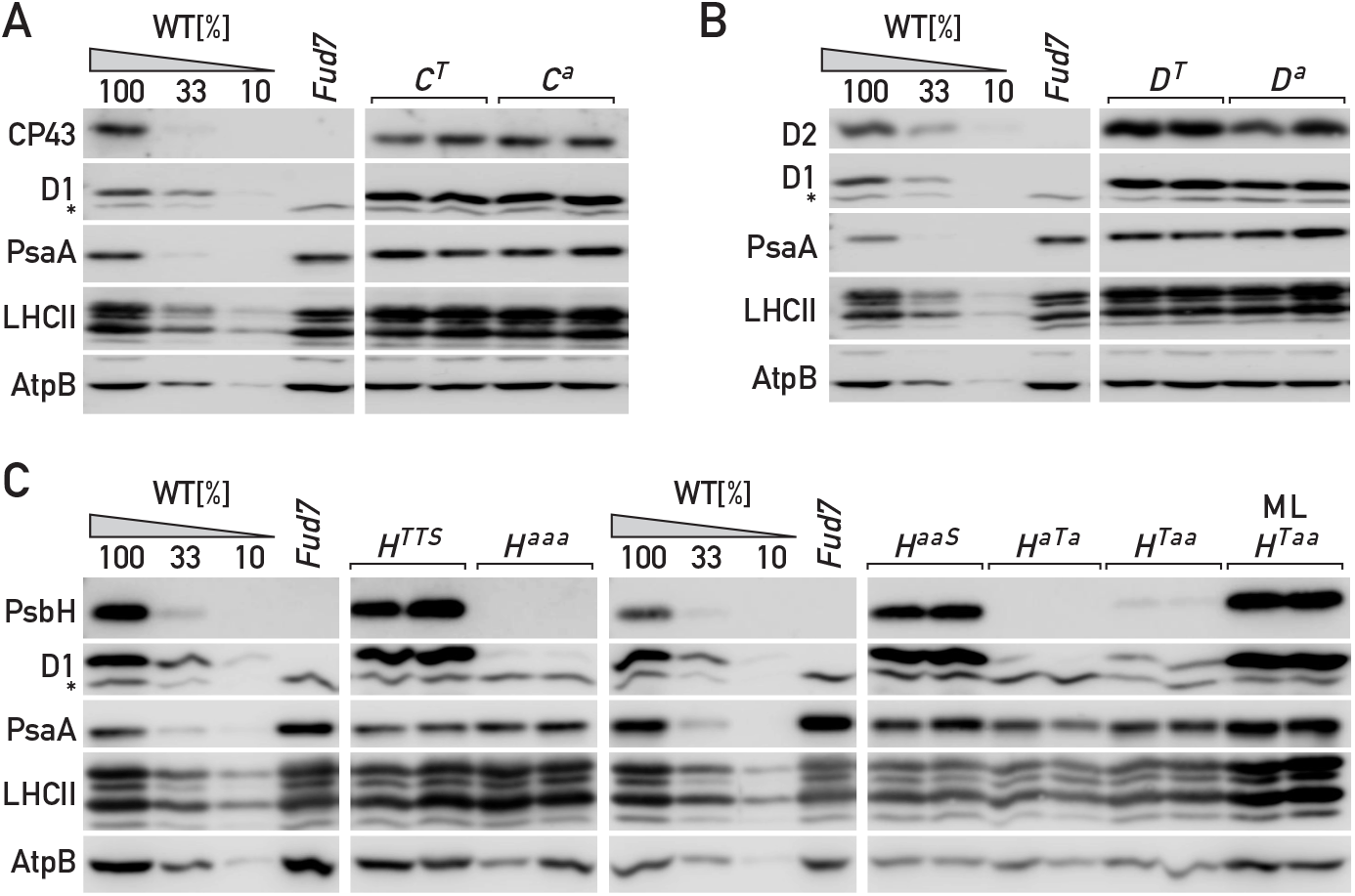
Immunoblot analysis of thylakoid proteins. Cells were grown in the dark and total protein extracts were analyzed by SDS-PAGE followed by immunodetection with antisera against subunits of photosystem II (CP43 (PsbC), D2 (psbD), PsbH or D1 (PsbA)), photosystem I (PsaA) and light-harvesting complex II (LhcbM5), as well as ATP synthase (AtpB) used as a loading control. Two independent lines are analyzed for each mutant genotype, and the immunoblots shown are representative of two independent experiments. Lower amounts of the wild-type extract (33%, 10%) are also loaded alongside for calibration of the immuno-detection. A non-specific band that cross-reacts with the D1 antiserum, but is still present in the deletion strain *Fud7*, is marked with an asterisk (*). (A) Alanine substitution in CP43 (PsbC) (B) Alanine substitution in D2 (psbD) (C) Alanine substitutions in PsbH. The two *H*^*Taa*^ strains were also grown at 60 μE m^−2^ s^−1^ (ML) for comparison to the dark-grown samples.

Thus the question arises whether in *H*^*Taa*^ the lower accumulation of PSII in the dark was due to a defect in its assembly or to an increase in its proteolytic turnover. To distinguish these alternatives, cultures of *H*^*Taa*^ and of *H*^*TTS*^ controls were grown in the light, so that they accumulated wild-type levels of PSII, and were then transferred to the dark in the presence or absence of inhibitors of chloroplast translation (lincomycin and chloramphenicol) which prevent *de novo* synthesis and assembly of PSII, for lack of its chloroplast-encoded subunits. The fate of PSII in the dark was monitored by chlorophyll fluorescence analysis and by immunoblotting. The maximum quantum yield of PSII (Fv/Fm) in the control remained stable over two days in the absence of the inhibitors, and slowly decreased in their presence (Fig 4A). The same diminution as in the inhibitor-treated control was observed in *H*^*Taa*^, in the presence or absence of inhibitors. Thus in the *H*^*Taa*^ mutant, PSII that was assembled in the light declined in the dark at the same rate as in the control with inhibitors. Monitoring the amount of the D1 subunit of PSII by immunoblotting gave similar results (Fig 4B,C). In the dark without the inhibitors, the control maintained constant levels of D1, while they gradually declined in the *H*^*Taa*^ mutant. In the presence of inhibitors, the amount of D1 dropped faster, and with similar rates in the two genotypes. These data strongly suggest that for *H*^*Taa*^, the reduced Fv/Fm and lower levels of PSII in the dark are due to a defect in *de novo* assembly of PSII and not to its destabilization.

**Figure 4.**
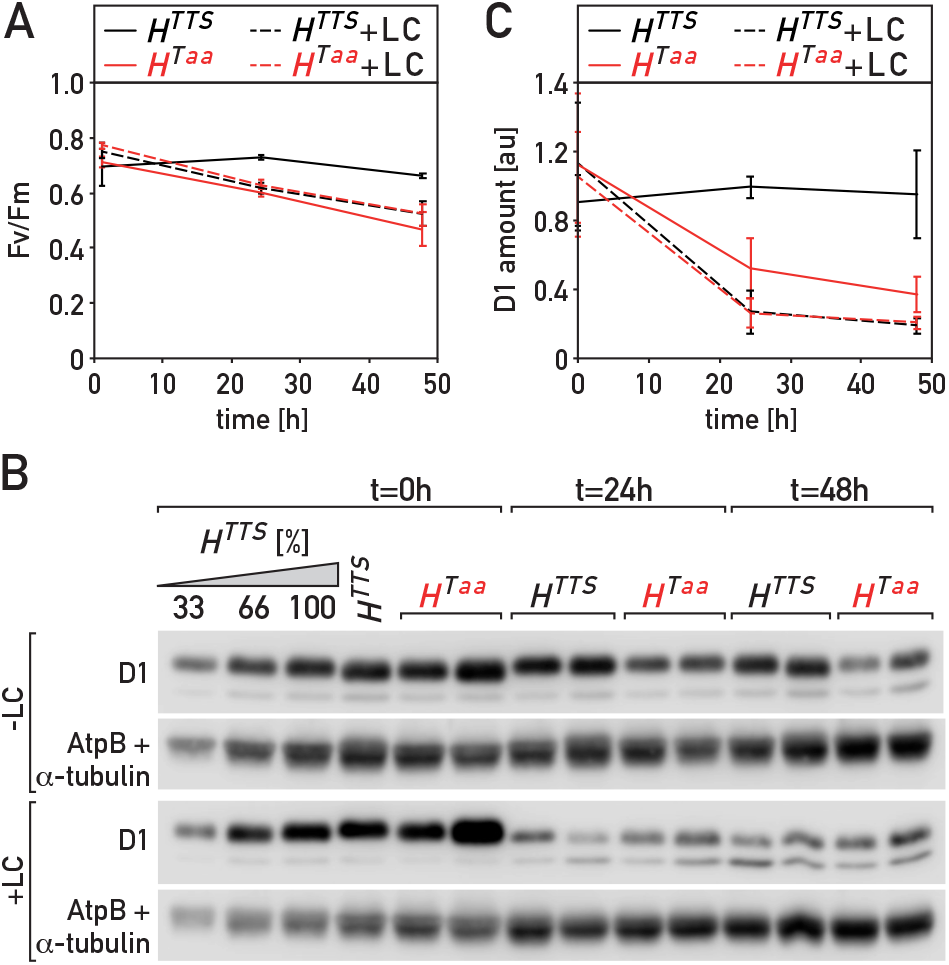
PSII turnover in the *H*^*Taa*^ mutant in the dark. Cells of the *H*^*Taa*^ mutant and the *H*^*TTS*^ control were grown in the light (60μE m^−2^ s^−1^) and then shifted to the dark for 48 hours in the presence (+LC) of the chloroplast translation inhibitors lincomycin and chloramphenicol or in their absence. (A) Maximum quantium yield of PSII (F_v_/F_m_). In the absence of inhibitors, *H*^*Taa*^ was significantly different from the control (Repeated Measures ANOVA, p<0.001). When inhibitors were applied, there was no significant difference with the control (p=0.355). (B) Total proteins separated by SDS-PAGE and immunodetected with antiserum against the D1 subunit of PSII. Antisera against AtpB and α-tubulin were used as a loading control. (C) Quantification of D1 in panel B (normalized to the loading controls). In the absence of inhibitors, *H*^*TTS*^ was significantly different from *H*^*Taa*^ (Repeated Measures ANOVA, p=0.047), especially at t=24h and t=48h (post-hoc test with Bonferroni’s correction, p=0.0083 and p=0.0137). There was no significant difference between *H*^*TTS*^ +LC and *H*^*Taa*^ + LC (p=0.837), or between *H*^*Taa*^ without inhibitors and either *H*^*TTS*^ +LC (p=0.837) or *H*^*Taa*^ + LC (p=0.229).

### Recovery from photo-inhibition is delayed in some of the phosphorylation mutants

PsbH harbors three major phosphorylation sites, and we sought to determine its degree of phosphorylation in the alanine substitution mutants using Phos-tag gel electrophoresis (Fig. 5). Phos-tag can be covalently incorporated into the polyacrylamide matrix and chelate divalent cations such as Zn^2+^, which in turn interact with phosphate groups (Kinoshita *et al.* 2009). Thus in Phos-tag gels, the migration of phosphorylated proteins is retarded compared to the non-phosphorylated form. After immunoblotting, the relative signals of the different forms denote the degree of phosphorylation of the protein. A wild-type culture was grown in the dark, and aliquots were either maintained in the dark (D) or transferred to low light (LL, 6 μE m^−2^ s^−1^), moderate light (ML, 60 μE m^−2^ s^−1^) or high light (HL, 250 μE m^−2^ s^−1^) for 1 hour. After Phos-tag PAGE of the protein extracts, two bands were detected. As a marker for the migration of the non-phosphorylated form, a sample was treated with a non-specific protein phosphatase prior to electrophoresis (λ phosphatase). The lower band migrated at the same size as a band detected in the sample treated with phosphatase, indicating that it corresponds to a non-phosphorylated form. The ratio of the signal between the higher band and the lower one indicated that the maximum degree of phosphorylation was observed in the culture transferred to moderate light. Those alanine substitution lines that do accumulate PSII were retained for the analysis (*H*^*Taa*^ and *H*^*aaS*^), using cultures that were grown in low light (to allow PSII accumulation in H^Taa^), dark-adapted for 1h and then transferred to moderate light for 1 hour to maximize PsbH phosphorylation. Compared to the strong phosphorylation of PsbH in the wild type and in the *H*^*TTS*^ control, phosphorylation was much lower in *H*^*Taa*^, and no phosphorylation was detected in the *H*^*aaS*^ mutant. Hence for further analysis we selected *H*^*aaS*^ because it has undetectable phosphorylation while retaining normal amounts of PSII, together with *C*^*a*^ and *D*^*a*^.

**Figure 5.**
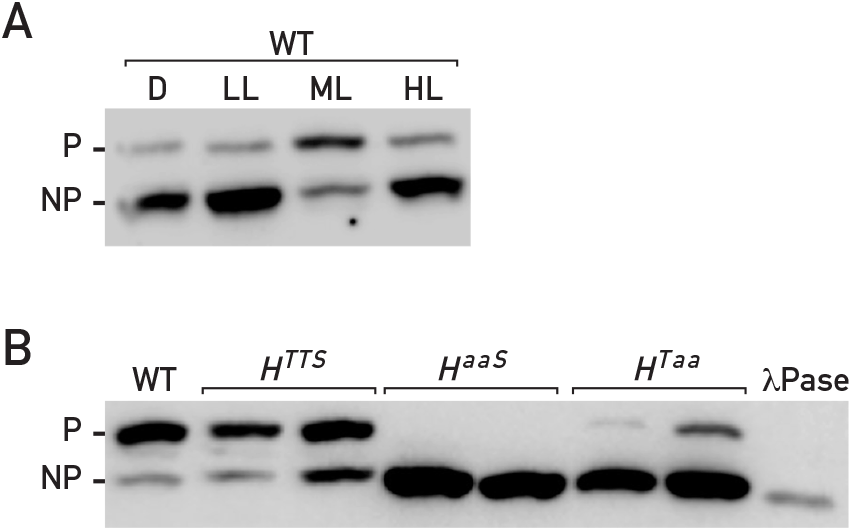
PhosTag analysis of protein phosphorylation. (A) Wild-type cells were grown in the dark and further incubated for 1h in the dark (D) or under different light intensities (LL: 6 μE m^−2^ s^−1^; ML: 60 μE m^−2^ s^−1^; HL: 250 μE m^−2^ s^−1^). Total protein extracts were analyzed by PhosTag-PAGE followed by immunoblotting with PsbH antiserum. P: phosphorylated; NP: non-phosphorylated. (B) Wild-type (WT), control (*H*^*TTS*^) and PsbH mutant lines (*H*^*aaS*^, *H*^*Taa*^) were grown in low light (to allow accumulation of PSII in H^Taa^), dark adapted for 1h and exposed for 1h to medium light (60 μE m^−2^ s^−1^). Total protein extracts were separated by PhosTag-PAGE followed by immunodetection with PsbH antiserum. Two independent lines are shown for each mutant genotype. As a reference for the migration of the fully de-phosphorylated protein, a total protein extract of *cw15* cells was treated *in vitro* with the non-specific protein phosphatase λ and loaded alongside (λ Pase). The immunoblots shown in A and B are representative of two independent experiments.

To determine whether protein phosphorylation plays a role in the repair cycle of the PSII, we analyzed photoinhibition in the alanine substitution mutants (Fig 6). Cells were grown in the dark, transferred to photoinhibitory light (1450 μE m^−2^ s^−1^) for 30 minutes and then allowed to recover in dim light (15 μE m^−2^ s^−1^) for 120 minutes. Photoinhibition was monitored by using chlorophyll fluorescence spectroscopy to determine the maximum quantum yield of PSII (Fv/Fm) at different time points. As a control for strong photoinhibition, we used the *ftsh* mutant which is deficient in a key protease of the PSII repair cycle (Malnoe, et al. 2014). At the end of the photoinhibitory treatment, Fv/Fm in the wild type dropped to 20% of its original value, and rapidly recovered in dim light to more than 80% of its initial value. In contrast Fv/Fm dropped to a very low level in *ftsh* (< 10% of the initial value) and failed to recover in dim light (Fig 6A). Photoinhibition and recovery were not significantly different in *D*^*a*^ and the *D*^*T*^ control (Fig 6B; p=0.835 in repeated measures ANOVA between the two genotypes), while a small delay was observed in the recovery of *C*^*a*^ compared to *C*^*T*^ (Fig 6C; p=0.0103), and a more significant delay for *H*^*aaS*^ compared to *H*^*TTS*^ (Fig 6D; p=0.0015).

**Figure 6.**
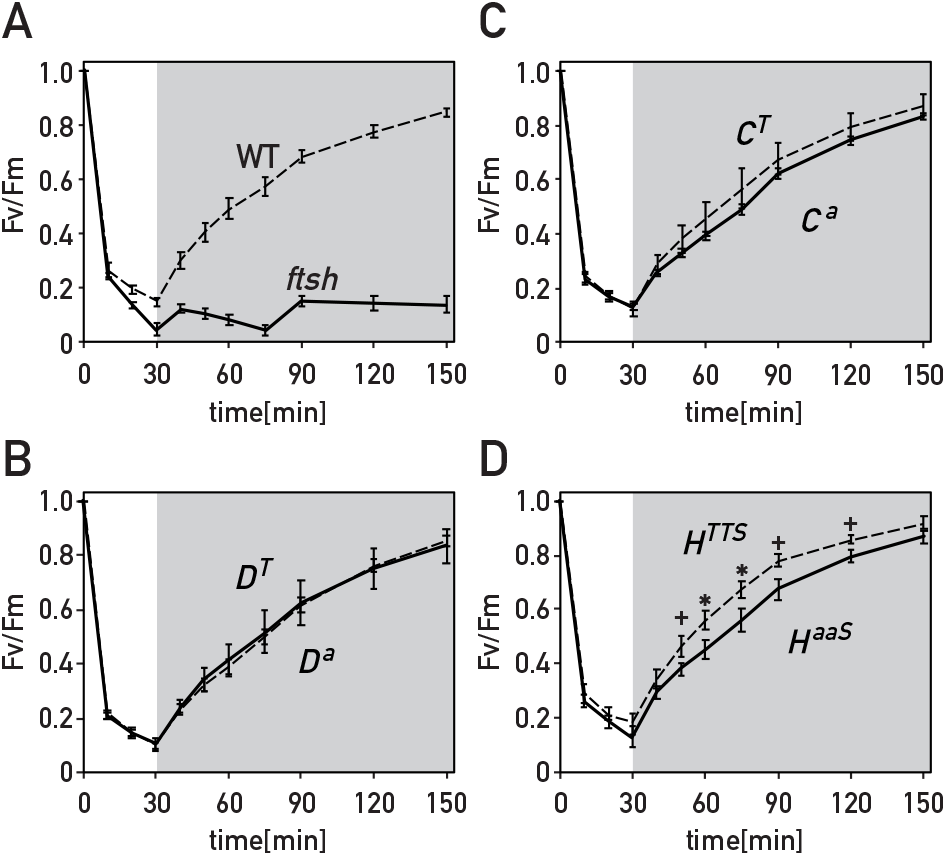
Kinetics of photoinhibition and recovery. Cells were grown in the dark, and then subjected to photoinhibition at 1450 μE m^−2^ s^−1^ for 30min followed by recovery at 15 μE m^−2^ s^−1^ for 120 min (gray shading). The maximum quantum yield of PSII (F_v_/F_m_) was determined at each time-point (after 2 minutes of dark adaptation) and normalized to its value before the treatment (t = 0). Each time-point is the average of four measurements on two independent lines of identical genotype (+/− SD). Significant differences at each time point were determined using Repeated Measures ANOVA and a post-hoc test with Bonferroni’s correction (+: p<0.005, *: p<0.001). (A) *ftsh1-1* (solid line) compared to the wild type (dashed line). (B-D) in each panel an alanine-substitution mutant (solid line) is compared to the corresponding control containing only a footprint of the recycled cassette at the respective locus (dashed line).

Although phosphorylation of CP43 and D2 was highly conserved during evolution, it was somewhat surprising that the individual alanine substitution mutants of these proteins had no strong phenotype. For PsbH, some mutants were strongly affected in the accumulation of PSII, while *H*^*aaS*^ had normal amounts but showed delayed recovery after photo-inhibition. Therefore to further investigate the role of phosphorylation in the repair of PSII, we analyzed lines with combined mutations in two of the target proteins (*C*^*a*^*D*^*a*^, *C*^*a*^*H*^*aaS*^ and *D*^*a*^*H*^*aaS*^), as well as lines with mutations in all three subunits (*C*^*a*^*D*^*a*^*H*^*aaS*^). The matching control lines carried the 483bp footprints at the respective loci. Immunoblotting with anti-phospho-threonine showed that phosphorylation of CP43 and PsbH is severely affected in the triple mutant genotype (Suppl. Fig S7), indicating as expected that the major phosphorylation sites of these proteins have been altered. Phosphorylation of D2 could not be assessed, as it migrated together with components of LHCII, but substitution of Thr2 with Ala was previously shown to abolish D2 phosphorylation (Fleischmann and Rochaix 1999). Growth tests showed that all lines were capable of photoautotrophic growth on minimal medium similar to the wild type and the controls (Suppl. Fig. S5). Immunoblotting with representative subunits indicated that the thylakoid complexes PSII, PSI, LHCII and ATP synthase are present in normal amounts in all the lines (Fig. 7). The parameters Fv/Fm and ΦPSII were also similar to those of the wild type in all the lines, indicating that PSII activity is not strongly affected (Fig. 8).

**Figure 7.**
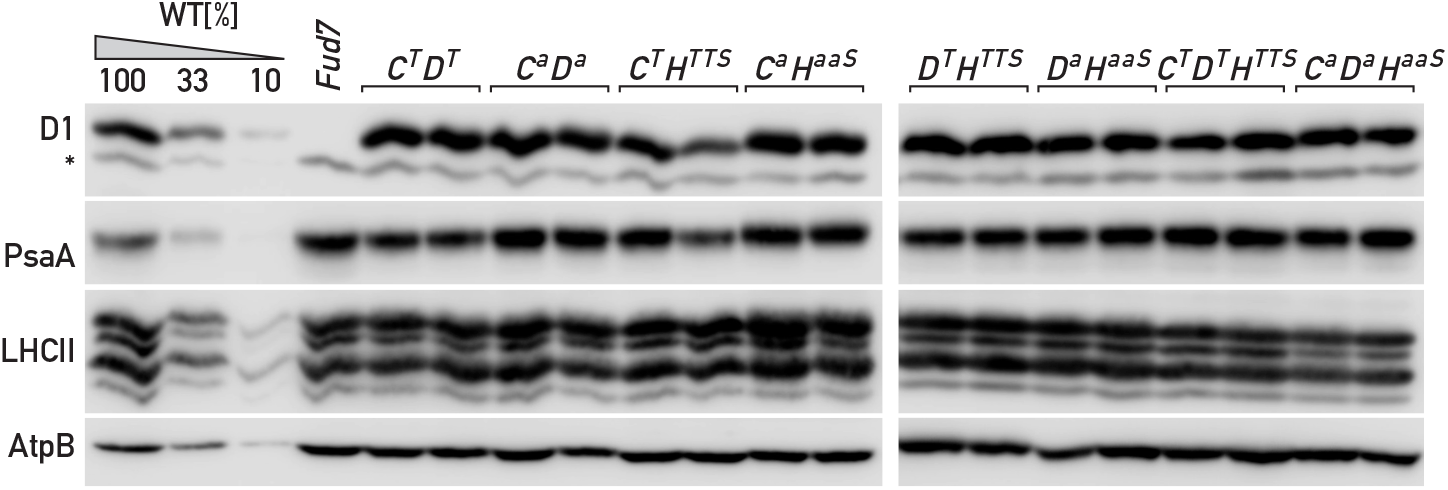
Immunoblot analysis of thylakoid proteins in multiple-site mutants. Cells were grown in the dark and analyzed as in Figure 3, with the antisera indicated on the left. Two independent lines are shown for each mutant genotype. Lower amounts of the wild-type extract (33%, 10%) are also loaded alongside for calibration of the immuno-detection. The immunoblots shown are representative of two independent experiments.

**Figure 8.**
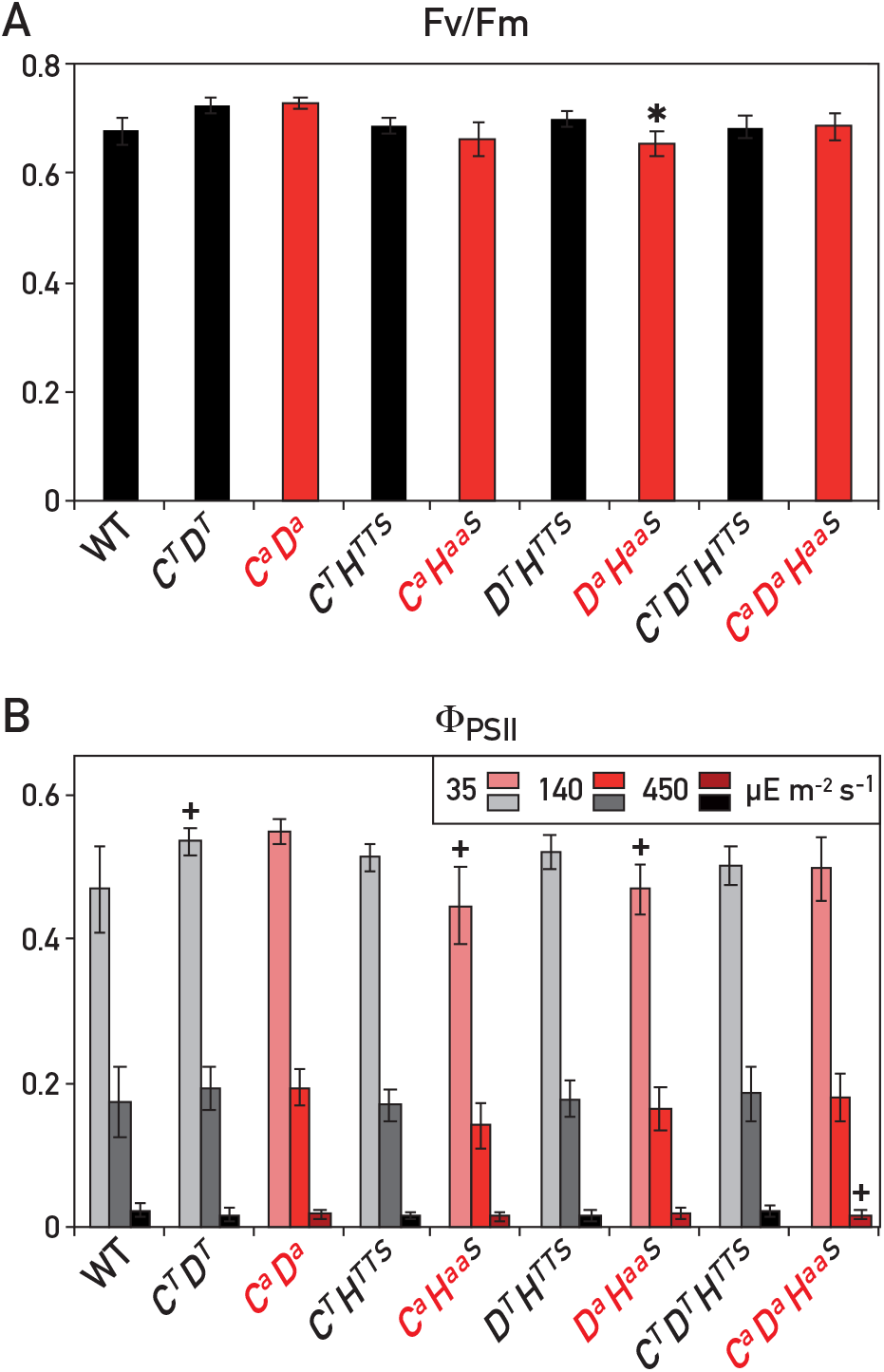
Chlorophyll fluorescence spectroscopy of mutants with alanine substitutions in two or three proteins. A. Maximum quantum yield of photosystem II (see Fig. 2) of strains with alanine substitutions in two or three proteins compared to the respective controls. B. Quantum yield of photosystem II (see Fig. 2). Each value is the average of three measurements on two independent lines of identical genotype (+/− SD). Significant differences in Student’s t-test between the control strains and the wild type, or between the mutants and their respective controls, are indicated (+: p<0.05; *: p<0.01).

In photoinhibition experiments, the double mutant *C*^*a*^*D*^*a*^ was not distinguishable from its control *C*^*T*^*D*^*T*^ (Fig 9A; p=0.77 in repeated measures ANOVA between the two genotypes). The double mutant *D*^*a*^*H*^*aaS*^ showed a small delay in recovery compared to *D*^*T*^*H*^*TTS*^ (Fig 9B; p=0.0005), similar to the delay observed in the *H*^*aaS*^ single mutant. However in the double mutant *C*^*a*^*H*^*aaS*^, a small increase in the rate of photoinhibition and a strong retardation of recovery were observed in comparison to the *C*^*T*^*H*^*TTS*^ control (Fig 9C; p=0.0001). To investigate whether these differences in *C*^*a*^*H*^*aaS*^ reflected changes in the sensitivity to photoinhibition or in the repair of photodamage, the kinetics of photoinhibition were compared in the presence or absence of chloroplast translation inhibitors (lincomycin and chloramphenicol, Suppl. Fig. S8). In the absence of chloroplast translation, damaged D1 subunits cannot be replaced by de-novo synthetized ones, so that the repair of PSII is deficient. The difference in the rate of photoinhibition between *C*^*a*^*H*^*aaS*^ and *C*^*T*^*H*^*TTS*^ which was apparent in the absence of inhibitor was clearly reduced in the presence of the translation inhibitors. Thus the difference observed in the absence of the inhibitors largely reflects repair activity occurring already during the photoinhibitory treatment in *C*^*T*^*H*^*TTS*^ but not in *C*^*a*^*H*^*aaS*^. These data indicate that it is mainly the repair of PSII rather than its photo-inactivation which is modified by the combined alanine substitutions. Surprisingly, in the triple mutant *C*^*a*^*D*^*a*^*H*^*aaS*^ the phenotype observed in double mutant *C*^*a*^*H*^*aaS*^ was no longer apparent, as the triple mutant showed rates of photoinhibition and recovery that were not significantly different from the control *C*^*T*^*D*^*T*^*H*^*TTS*^ (p=0.854 in ANOVA between the two genotypes).

**Figure 9.**
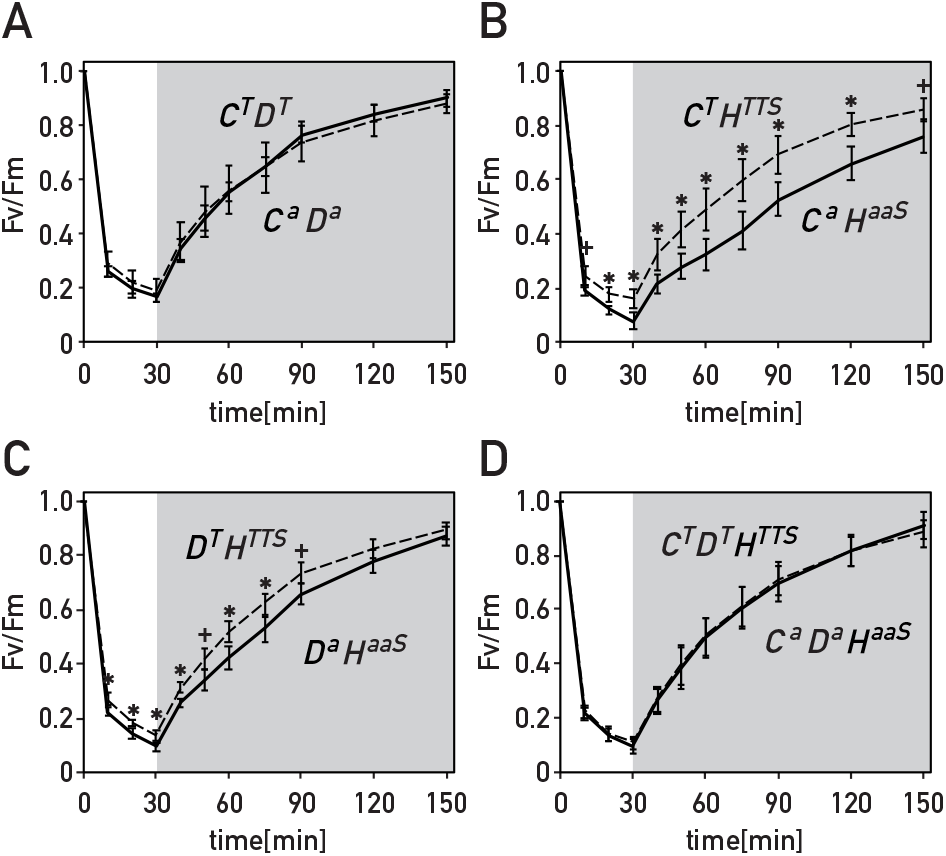
Kinetics of photoinhibition and recovery of mutants with alanine substitutions in two or three proteins. In each panel a mutant with alanine substitutions in two or three proteins (solid line) is compared to the corresponding control containing only the footprints of the recycled cassette at the respective loci (dashed line). Cells were grown, treated and analyzed as in Fig. 6.

## DISCUSSION

In the interpretation of the phenotypes of the alanine substitution mutants, it is important to consider whether they are due to the lack of phosphorylation, or to a structural change imparted on the protein by the substitution of a hydroxylated threonine or serine residue with a hydrophobic alanine. Phosphorylation of the PSII core subunits CP43, D2 and PsbH occurs at or near the N-terminus of the mature proteins. This is a region of low structural constraint that is not resolved in the cryo-EM structures of PSII (Table 1), suggesting that the mutations should not have a major effect on the structure of PSII. Indeed for CP43 and D2 the alanine substitutions must have minimal deleterious structural consequences, because the mutants *C*^*a*^ and *D*^*a*^ do not show any significant phenotype. However for PsbH, it should be considered whether the alanine substitutions could have damaging effects on the structure of PSII. From the normal PSII accumulation and quantum yield of *H*^*aaS*^ and *H*^*Taa*^ grown in the light, it appears that none of the three residues is individually essential. Furthermore these same two mutants show that the presence of two additional hydrophobic alanine residues can be tolerated in the N-terminal domain, which normally already contains one alanine. That the presence of three supernumerary alanines could be structurally detrimental remains a possibility which cannot be excluded since *H*^*aaa*^ accumulates only low levels of PSII proteins. For these reasons, in our analysis of photoinhibition we used only lines that accumulated normal levels of fully active PSII, so that the phenotypes we observed could be ascribed to the lack of phosphorylation rather than to structural defects.

Although phosphorylation of the PSII core subunits CP43 and D2 at their N-terminus was conserved during evolution (reviewed by (Grieco, et al. 2016)), the *C*^*a*^ and *D*^*a*^ mutants did not exhibit any significant phenotype in their photoautotrophic growth or quantum yield of PSII, and C^a^ presented only a minor susceptibility to photoinhibition and subsequent recovery. Similar observations were previously reported for the *D*^*a*^ mutant (Andronis, et al. 1998, Fleischmann and Rochaix 1999). Thus phosphorylation of CP43 or D2 is not essential for photosynthesis, at least under normal laboratory conditions. In contrast some alanine substitutions of the phosphorylation sites in PsbH had much more drastic effects on the stable accumulation of PSII and affected recovery from photoinhibition. Simultaneous mutation of all three sites (Thr3, Thr5 and Ser6) led to a nearly complete loss of PSII in the *H*^*aaa*^ mutant, similar to the deletion mutant Δ*psbH*. In H^*aTa*^, the presence of only Thr5 was not sufficient to fully rescue this phenotype, indicating a crucial role for Thr3 and Ser6 that cannot be compensated by Thr5 alone. However the presence of only Ser6 (*H*^*aaS*^) was sufficient to restore PSII accumulation and activity to wild-type levels. This is consistent with the previous report that a mutant corresponding to *H*^*aTS*^ had no major photosynthetic phenotype (O’Connor, et al. 1998). Furthermore the presence of Thr3 (*H*^*Taa*^) was also sufficient during growth in the light, suggesting that Thr3 and Ser6 show some redundancy in their function. However during growth of *H*^*Taa*^ in the dark the accumulation of PSII was only partial. When light-grown *H*^*Taa*^, containing normal amounts of PSII, was transferred to the dark in the presence of protein synthesis inhibitors, the half-life of PSII was similar in the mutant and in the control *H*^*TTS*^. Thus the reduced accumulation of PSII in *H*^*Taa*^ grown in the dark seems to be due to a difference in assembly rather than in degradation. While a defect in translation initation cannot be ruled out, this seems unlikely since the mutation affects a nucleotide which is 13 bases downstream of the AUG start codon. Hence the differences in the phenotypes of *H*^*Taa*^ in the light versus the dark point to a requirement for light in the assembly of PSII in this mutant, through an unknown mechanism which it would be interesting to unravel in future investigations. Taken together, our results point to a central role of PsbH and its amino-terminal domain in the assembly or stability of PSII.

The mutant retaining only Ser6 (*H*^*aaS*^) is competent for assembly and maintenance of active PSII, while the mutant further lacking this serine (*H*^*aaa*^) is not. However using Phos-tag gel electrophoresis and immunoblotting, we could not detect any phosphorylation of PsbH in *H*^*aaS*^. We can offer two hypotheses to possibly explain these observations. One is that phosphorylation of Ser6 could depend on the prior phosphorylation of Thr3 or Thr5, consistent with the detection of only di-phosphorylated peptides by mass spectrometry (Lemeille, et al. 2010, Wang, et al. 2014). This would imply that phosphorylation of PsbH would not be strictly required for assembly and stable PSII accumulation. The other hypothesis is that Ser6 might be only very transiently phosphorylated during PSII assembly, so that phosphorylation of this residue is not detectable at steady state. In fact the latter hypothesis might be extended to the D1 subunit of Chlamydomonas, which was not detectably phosphorylated after ^32^P labelling (de Vitry, et al. 1991, Delepelaire 1984), although its phosphorylation could be observed by mass spectrometry (Turkina, et al. 2006, Wang, et al. 2014). In plants it has been proposed that D1 undergoes a phosphorylation and de-phosphorylation sequence during PSII repair (Baena-Gonzalez, et al. 1999). Thus it cannot be ruled out that in Chlamydomonas the phosphorylation of D1 might be too low and transient to allow its detection by ^32^P labelling at steady state in the presence of other strongly phosphorylated proteins.

Sensitivity to photo-inhibition and subsequent recovery was not measurably affected in the individual mutant of D2 (*D*^*a*^), indicating that phosphorylation of this residue is not essential in the repair cycle of PSII, while the mutant of CP43 (*C*^*a*^) showed a small delay in recovery. The recovery from photo-inhibition was affected in the *H*^*aaS*^ mutant of PsbH, and most strongly so in the *C*^*a*^*H*^*aaS*^ line with the combination of mutations in PsbH and CP43. These data indicate that PsbH phosphorylation may play an important role in the repair of PSII after photo-inhibition. Surprisingly, the phenotype of *C*^*a*^*H*^*aaS*^ was suppressed in the mutant combination *C*^*a*^*D*^*a*^*H*^*aaS*^ with the additional mutation in D2. Likewise the mild phenotype of *C*^*a*^ was no longer apparent in the double mutant *C*^*a*^*D*^*a*^. However the phenotype of *D*^*a*^*H*^*aaS*^ was not different from *H*^*aaS*^, suggesting that the suppressive effect of *D*^*a*^ is exerted only in the presence of the *C*^*a*^ mutation. Taken together all these data suggest that phosphorylation of PSII, and in particular of PsbH, has an important role in its repair cycle, but that this is not simply an additive effect. There are several ways in which phosphorylation of the subunits of PSII could be involved in the repair cycle: indirectly through changes in thylakoid architecture, or more directly through changes in the stability of PSII supercomplexes, or through altered molecular recognition by some of the numerous actors of repair (Jarvi, et al. 2015, Komenda, et al. 2012). The photo-inhibition phenotypes of the mutant lines are discussed below in the alternative perspectives of an indirect or direct role of PSII phosphorylation.

Thylakoid stacking depends on a balance of attractive and repulsive forces that is influenced by protein phosphorylation and the negative charges that this entails, as well as by the concentrations of monovalent and divalent cations (Puthiyaveetil, et al. 2017). It has been proposed that a minimal threshold of phosphorylation is required for thylakoid stacking (Fristedt *et al.* 2010), but that strong phosphorylation increases repulsive electrostatic forces and reduces stacking interactions. In turn thylakoid stacking defines domains of the thylakoid network where different steps of the repair cycle are thought to occur, namely grana, grana margins and stromal lamellae. Although in Chlamydomonas grana are not as distinct, the thylakoid network is also organized in similar domains (Engel *et al.* 2015, Vallon *et al.* 1985). The PSII complexes and supercomplexes are usually found within the tightly appressed membranes of the grana, with little access to players of the repair cycle such as proteases located in the grana margins and the translation apparatus at the stromal lamellae (Puthiyaveetil, et al. 2014a). Thus phosphorylation of PSII may facilitate repair by loosening the grana, allowing access of damaged PSII to the grana margins and facilitating contact with the repair machinery. In this perspective, the photoinhibition phenotypes of the mutants might be explained (i) by insufficient phosphorylation for electrostatic repulsion and grana unstacking in *C*^*a*^*H*^*aaS*^, hence its defect in PSII repair, but (ii) by phosphorylation below the required threshold in *C*^*a*^*D*^*a*^*H*^*aaS*^, with reduced stacking and easier access of the repair machinery.

Repair of photo-damaged PSII is thought to initiate with the disassembly of PSII-LHCII complexes, to facilitate diffusion of PSII to the grana margins and facilitate subsequent steps, such as access of the proteases, selective dissociation of some subunits of PSII, and replacement with intact subunits (Jarvi, et al. 2015, Komenda, et al. 2012). The dismantling of the PSII-LHCII supercomplexes may be facilitated by protein phosphorylation resulting in repulsive electrostatic forces between phosphorylated PSII and LHCII (Puthiyaveetil, et al. 2017, Tikkanen, et al. 2008), while monomerization of PSII was proposed to be inhibited by phosphorylation (Kruse *et al.* 1997). The repair of PSII involves a further partial disassembly of PSII monomers with the transient removal of CP43, which might be aided by its de-phosphorylation (Baena-Gonzalez, et al. 1999). Furthermore in plants there is biochemical evidence that the degradation of D1 and D2 after photo-damage is impeded by their phosphorylation, so that the repair cycle requires a step of de-phosphorylation (Ebbert and Godde 1996, Koivuniemi, et al. 1995). There is also genetic evidence that phosphorylation by STN8 protects PSII from proteolytic degradation (Kato and Sakamoto 2014). In this perspective, phosphorylation-induced dissociation of the PSII supercomplexes in *C*^*a*^*H*^*aaS*^ might be insufficient for efficient repair, but this defect could be counteracted and thus suppressed by the *D*^*a*^ substitution in *C*^*a*^*D*^*a*^*H*^*aaS*^, as the lack of D2 phosphorylation would facilitate recognition by the proteolytic machinery.

As discussed above, both direct and indirect effects of phosphorylation can be invoked to rationalize the phenotypes of the phosphorylation-site mutants. These different hypotheses are not mutually exclusive. Whatever the mechanisms, our observations in Chlamydomonas suggest that changes in PSII phosphorylation may explain at least in part the defects in PSII repair that were observed in the *stn8* and *pbcp* mutants of Arabidopsis (Kato and Sakamoto 2014, Puthiyaveetil, et al. 2014b, Tikkanen, et al. 2008). Furthermore our results indicate that phosphorylation of PSII should not be considered only in bulk, and that the phosphorylation of its different subunits may have distinct functions. In particular our data reveal a prominent role for phosphorylation of PsbH in PSII assembly and in its repair after photo-inhibition.

## MATERIAL AND METHODS

### Strains and growth conditions

*Chlamydomonas reinhardtii* wild type strain 2D+, (a derivative of 137C) was used for chloroplast transformation. *C. reinhardtii* mutants were previously described, *Fud7* (Bennoun, et al. 1986), and *ftsh 1.1 Su*^−^ (Malnoe, et al. 2014).

Cells were grown in Tris-acetate-phosphate medium (TAP) in the dark (unless indicated otherwise) or in High Salt Minimal medium (HSM) under 60 μE m^−2^ s^−1^ white light from fluorescent tubes (Gorman and Levine 1965, Sueoka 1960). They were collected for analysis during the exponential growth phase (2.10^6^ cells.mL^−1^). When stated, inhibitors of the chloroplast translation machinery, lincomycin and chloramphenicol, were added to the culture at a concentration of 500 and 100 μg mL^−1^ respectively.

### DNA constructs and site-directed mutagenesis

Plasmid pAR1-*psbH*/*aadA* was obtained by ligation of the recyclable *aadA* selection cassette (as a *MluI*-*MluI* fragment) (Fischer, et al. 1996, Goldschmidt-Clermont 1991) into the *MluI* site of plasmid p41 (which contains the 3.2 kb *BamHI*-*ScaI* fragment of *psbH*) (Loizeau *et al.* 2014).

pAR14-*psbC*/*aadA* was obtained by insertion of a recyclable *aadA* selection cassette (as a *MluI*-*MluI* fragment) into a plasmid containing a 3.7 kb XbaI-SalI *psbC* fragment, at an engineered *MluI* restriction site 733 bp upstream of the GTG start codon of *psbC*. This site turned out to be located in the 3’UTR of the *ycf2* gene (Cavaiuolo *et al.* 2017), but after excision of the *aadA* marker the control transformants with the residual footprint (*C*^*T*^) had no apparent phenotype.

pAR8-*psbD*/*aadA* was obtained by replacing the non-recyclable *aadA* cassette *ClaI*-*SpeI* fragment by a recyclable *aadA* cassette (*NheI*-*ClaI*) in a plasmid containing the 2.9 kb *EcoRI* fragment of *psbD* (Fleischmann and Rochaix 1999).

A panel of derivatives of these three plasmids with mutations at several phosphorylation sites was created using PCR with mutated primers and Gibson assembly (Gibson 2011) (table III.S1).

### Chloroplast transformation

Transformation was performed using the biolistic method with a Helium gene gun. Transformants were selected on TAP-agar plates in the presence of 150 μg.mL^−1^ spectinomycin and repeatedly sub-cultured on the same medium. Homoplasmy for the presence of the *aadA* cassette was checked by PCR amplification of the region surrounding the insertion site. The presence of the desired mutations was confirmed by PCR with one primer selective for the mutation and the other in a nearby flanking region. Homoplasmy of the mutation was determined by a similar PCR with a primer specific for the wild-type sequence (Suppl. Fig S2). Alternatively, the presence of the mutation and its homoplasmy were deduced from bulk sequencing of a PCR amplicon surrounding the mutation site (Suppl. Table S2 and Suppl. Fig S3).

After reaching homoplasmy, the strains were repeatedly sub-cultured on spectinomycin-free TAP-agar plates. In the absence of selection pressure, the *aadA* cassette is removed by homologous recombination between the two flanking direct repeats, leaving only one repeat as a footprint in place of the cassette. Homoplasmic loss of the *aadA* cassette was checked by PCR amplification of the region flanking the insertion site and of an internal fragment of the cassette. Strains carrying only the footprint were used in the subsequent analysis, or in further rounds of transformation to obtain mutants in multiple genes.

### Chlorophyll fluorescence analysis

Chlorophyll fluorescence was monitored (Maxwell and Johnson 2000) using a green LED light source and a CCD camera (Speedzen system, JBeamBio, France). Maximum fluorescence *Fm* was measured with a 250 ms saturing flash.

ɸPSII measurements were performed on TAP-grown cells washed twice with High Salt Minimum medium (HSM) and allowed to recover for 1h at 6 μE m^−1^ s^−1^. Aliquots of the culture were adapted for 5 min to the specified light intensities, allowing ɸPSII values to stabilize before the measurement.

### Protein extraction and immunoblot analysis

Cells were resuspended in 100mM Tris pH 6.8, 2% SDS, 1x protease inhibitors (Sigma P8849) and incubated at 37°C for 30 min. Proteins were separated by SDS-PAGE in 10 or 12% SDS-polyacrylamide gels and transferred onto nitrocellulose membranes (0.45μm BioRad or 0.2μm Whatman) in a wet blotting apparatus. Loading of all gels was on an equal protein basis.

Antisera were used at the following dilutions: D1, CP43 and ATP synthase β-subunit (Agrisera) at 1:10,000 dilution, D2 (gift of J.-D. Rochaix) at 1:2,000, PsbH at a 1:5,000, PsaA (gift of Kevin Redding) at 1:50,000, and LhcbM5 (gift of Yuichiro Takahashi) at 1:20,000 dilution.

### Thylakoid membrane extraction

20mL of a culture exposed 1h to ML were sonicated in ice (3 times 30s, 30s rest in between, duty cycle 50%, output 50, Brandson Sonifier 450) in the presence of 1x protease inhibitiors (Sigma P8849) and 10mM sodium fluoride. Unbroken cells were pelleted at 3000g for 10min. The supernatant was transferred to a new tube and thylakoid membranes were pelleted at 75 000g for 30min at 4°C. Membranes were stocked in storage buffer (BisTris HCl 50mM pH 7, 750mM ACA, 20% glycerol).

### Protein phosphorylation analysis

For protein phosphorylation analysis, culture samples were quickly quenched in 4 volumes of cold acetone (−20°C) in order to preserve the relevant phosphorylation state. After precipitation at −20°C for at least 16 hrs, proteins were recovered by centrifugation and solubilized in 100mM Tris pH 7.5, 2% LDS, 1X protease inhibitors (ROCHE 04 693 132 01) and 10mM sodium fluoride for 30 min at 37°C. Samples were desalted on Zeba Spin columns following manufacturer instructions (7K MWCO, Thermo Scientific).

Proteins were separated by SDS-PAGE in 15% polyacrylamide gels containing 350mM Bis-Tris pH 6.8. Running buffer was prepared freshly (50mM MOPS, 50mM Tris, 0.1% SDS and 5mM sodium bisulfite). Transfer to nitrocellulose membranes was described above. Antiserum against phospho-Threonine (Zymed) at a 1:3,000 dilution was used for immuno-detection.

Separation of phosphorylated forms of proteins by LDS-PAGE in 10% polyacrylamide gels was performed using the Zn/Phos-tag method (Wako)(Kinoshita, et al. 2009). The resolving gel was supplemented with 100μM ZnNO_3_ and 50μM Phos-tag. Running buffer was prepared freshly and contained 50mM MES, 50mM Tris, 0.1% SDS and 5mM sodium bisulfite. Transfer and immunodetection were performed as described above.

### Photoinhibition

Cells in exponential phase (2-3.10^6^ cells.mL^−1^) were adjusted to 2.10^6^ cells.mL^−1^ and 20 mL were transferred to 100mL Erlenmeyer flasks. Photoinhibition was performed on an inverted white LED panel (JBealBio, France) mounted on a rotary shaker during 30 min at 1450 μE m^−2^ s^−1^. Recovery was performed at 15 μE m^−2^ s^−1^ for 2h. When needed, inhibitors of the chloroplast translation machinery, lincomycin and chloramphenicol, were added to the culture at a concentration of 500 and 100 μg.mL^−1^ 10 min before photoinhibition.

## ACKNOWLEDGEMENTS

This work was supported by the EU Marie Curie Initial Training Network (ITN) CALIPSO (GA ITN-2013 607 607) and by the the Swiss National Science Foundation (SNF 31003A_146300). We thank Jean-David Rochaix, Paolo Longoni, Geoffrey Fucile, Federica Cariti, Roman Ulm and Michael Hippler for scientific discussions, Emilie Demarsy for comments on the manuscript, José Manuel Nunes for help with the statistical analysis and Nicolas Roggli for preparing the figures.

## SUPPORTING INFORMATION

Table S1. Vectors and oligonucleotides used for site-directed mutagenesis.

Table S2. Primers used for genotyping of the transformants.

Figure S1. Construction of site-directed mutant vectors by Gibson assembly.

Figure S2. Genotyping of the transformants.

Figure S3. Sequencing profiles of wild-type and mutant DNA mixtures.

Figure S4. Pedigree of the mutant strains.

Figure S5. Growth tests.

Figure S6. Maximum quantum yield of PSII in PsbH mutants after growth in the light.

Figure S7. Immunoblot analysis of protein phosphorylation.

Figure S8. Effect of translation inhibitors on photoinhibition.

