## Supplemental Figures for "A key role for phosphorylation of PsbH in the biogenesis and repair of photosystem II in Chlamydomonas"

**Table S1. Vectors and oligonucleotides used for site-directed mutagenesis.**

The plasmids used for transformation were modified using Gibson assembly to introduce the desired phosphorylation-site mutations (see Fig S1). The table lists the transformation vector (plasmid vector), the restriction enzymes used for its digestion, and the oligonucleotides used for PCR amplification of the two overlapping fragments used to replace the excised sequence, with the mutated nucleotides shown in red and lowercase.

| **Mutant (shorthand)** | **Plasmid vector** | **Restriction enzymes** | **Oligonucleotide primers**<br>**A/B**<br>**C/D** |
|---|---|---|---|
| *PsbC T3A* *($C^{a}$)* | pAR14 | PacI-EcoRI | AAAAGTGcTTCCACTTTGCATTACCTCC /<br>CAGTTGTCATGTTAATGGAGCTTACTTTACC |
| | | | CGGAGGTAATGCAAAGTGGAAgCACTTTTTAATGG /<br>TAAACCTGTAGGACCTTGTGCAGAAGC |
| *PsbD T2A* *($D^{a}$)* | pAR8 | ClaI-PvuII | TGATATGTACCGATCGCAATTGcCATTGCGTGTATCTCC /<br>CCCGGTACCCAGCTTTTGTTCATCG |
| | | | gCAATTGCGATCGGTACATATCAAGAGAAACGCAC /<br>CACCGAAACTAGGTGCAAAGAACCAACCTG |
| *PsbH T3A T5A S6A* *($H^{aaa}$)* | pAR1 | PacI-NheI | AATCAATTATGGCAgCAGGAgCTgCTAAAGCTAAACC /<br>GCGGTGCCACTGCCGAATATAAAT |
| | | | cAGcTCCTGcTGCCATAATTGATTAAATGAATTAAGC /<br>GCAGTAGCCATTCATGTCACTTTGAAAC |
| *PsbH T3A T5A* *($H^{aaS}$)* | pAR1 | PacI-NheI | AATCAATTATGGCAgCAGGAgCTTCTAAAGCTAAACC /<br>GCGGTGCCACTGCCGAATATAAAT |
| | | | AAGcTCCTGcTGCCATAATTGATTAAATGAATTAAGC /<br>GCAGTAGCCATTCATGTCACTTTGAAAC |
| *PsbH T3A S6A* *($H^{aTa}$)* | pAR1 | PacI-NheI | AATCAATTATGGCAgCAGGAACTgCTAAAGCTAAACC /<br>GCGGTGCCACTGCCGAATATAAAT |
| | | | cAGTTCCTGcTGCCATAATTGATTAAATGAATTAAGC /<br>GCAGTAGCCATTCATGTCACTTTGAAAC |
| *PsbH T3T T5A S6A* *($H^{Taa}$)* | pAR1 | PacI-NheI | AATCAATTATGGCAACAGGAgCTgCTAAAGCTAAACC /<br>GCGGTGCCACTGCCGAATATAAAT |
| | | | cAGcTCCTGTTGCCATAATTGATTAAATGAATTAAGC /<br>GCAGTAGCCATTCATGTCACTTTGAAAC |

**Table S2. Primers used for genotyping of the transformants.**

The primers were used for genotyping as described in Figures S2. "N.S." indicates that the amplification was not specific enough to discriminate the wild-type and mutant sequences. Note that the same mutation-specific primer "I" (see Fig S2) was used for genotyping of $H^{aaa}$ and $H^{Taa}$.

| **Mutant** | **PCR primers *aadA* cassette E/F** | **PCR primer insertion site G/H** | **PCR primers mutant I/J (annealing T°)** | **PCR primers wild type K/L (annealing T°)** | **Primer for sequencing M** |
|---|---|---|---|---|---|
| *PsbC T3A* (*$C^{a}$*) | TTAGCTGGATAACGCCACGGAATG / ATGGCAATGCGTACTCCAGAAGAAC | CCAAAGCGTCGTCGTTGGTTAC / CTAAAGTTTATCGAACTTGCCACCATG | TTCCATTAAAAAGTGc / GTTAATGGAGCTTACTTTACC (51°C) | N.S. | CAGTTGTCATGTTAATGGAGCTTACTTTACC |
| *PsbD T2A* (*$D^{a}$*) | TTAGCTGGATAACGCCACGGAATG / ATGGCAATGCGTACTCCAGAAGAAC | TGGGACTAGAACTGCTTTGTGCATAG / CTTCGGGACGTCCTTACGGG | N.S. | N.S. | AATATAATAATTGTGATGACTATGC |
| *PsbH T3A T5A S6A* (*$H^{aaa}$*) | TTAGCTGGATAACGCCACGGAATG / ATGGCAATGCGTACTCCAGAAGAAC | CAGGCAACTGCCACTGACG / TGACGTTTCTATGAGTTGGGAAACTTTAGC | TCATTTAATCAATTATGGCAgCAGGAgCTg / GCGGTGCCACTGCCGAATATAAAT (56°C) | GGCAACAGGAACTT / AATATGGTTGAGTTGC (54°C) | GCGGTGCCACTGCCGAATATAAAT |
| *PsbH T3A T5A* (*$H^{aaS}$*) | TTAGCTGGATAACGCCACGGAATG / ATGGCAATGCGTACTCCAGAAGAAC | CAGGCAACTGCCACTGACG / TGACGTTTCTATGAGTTGGGAAACTTTAGC | TTTAGAAGcTCCTGc / CCATTCATGTCACTTTG (54°C) | TTATGGCAACAGGAA / GCGGTGCCACTGCCGAATATAAAT (55°C) | GCGGTGCCACTGCCGAATATAAAT |
| *PsbH T3A S6A* (*$H^{aTa}$*) | TTAGCTGGATAACGCCACGGAATG / ATGGCAATGCGTACTCCAGAAGAAC | CAGGCAACTGCCACTGACG / TGACGTTTCTATGAGTTGGGAAACTTTAGC | N.S. | N.S. | GCGGTGCCACTGCCGAATATAAAT |
| *PsbH T5A S6A* (*$H^{Taa}$*) | TTAGCTGGATAACGCCACGGAATG / ATGGCAATGCGTACTCCAGAAGAAC | CAGGCAACTGCCACTGACG / TGACGTTTCTATGAGTTGGGAAACTTTAGC | TCATTTAATCAATTATGGCAgCAGGAgCTg / GCGGTGCCACTGCCGAATATAAAT (40°C) | GGCAACAGGAACTT / AATATGGTTGAGTTGC (46°C) | GCGGTGCCACTGCCGAATATAAAT |

#### Figure S1

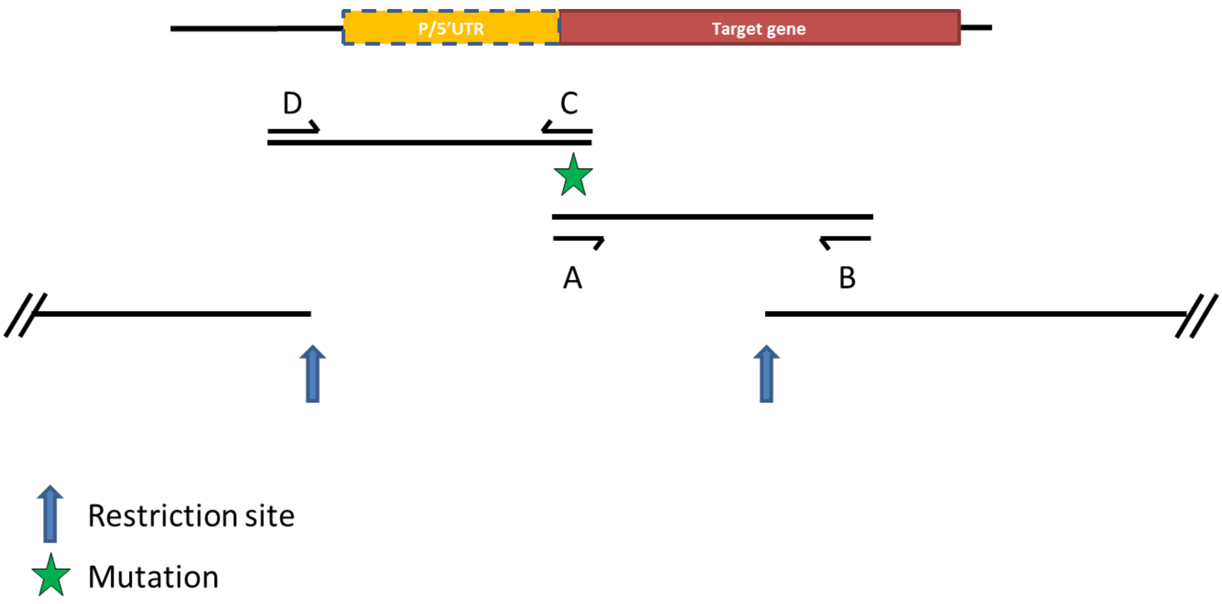


**Figure S1. Construction of site-directed mutant vectors by Gibson assembly**
Two overlapping PCR fragments are amplified, with the desired mutation (star) carried by primers A and C. The vector is digested with restriction enzymes (blue arrows) and a Gibson assembly is performed to replace the missing fragment with the two PCR fragments.

### Figure S2

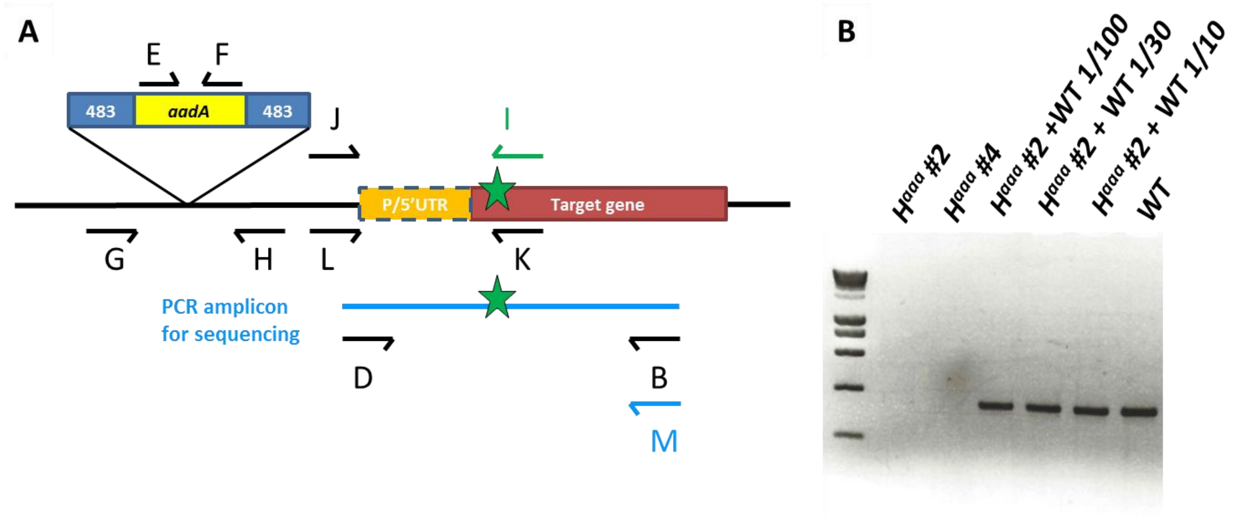


**Figure S2. Genotyping of the transformants.**

**A**. The schematic representation (not to scale) shows the insertion site of the excisable *aadA* cassette flanked by 483 bp direct repeats, as well as the promoter, 5'UTR (P/5'UTR) and coding sequence of the target gene, with the position of the site-directed mutation indicated by a green star. The primers used for genotyping by PCR are indicated as arrows labeled with capital letters, with the primer shown in green matching at its 3'end the mutant sequence (green star). The sequences of all primers are shown in Table S2. The presence or absence of the *aadA* cassette was verified by PCR amplification using the oligonucleotide primers E and F, listed in Table S2. Homoplasmy was deduced from the absence of amplification of the wild type sequence (primers G and H). The presence and homoplasmy of the desired mutation (marked with a green star) was verified by PCR amplification with primers matching the mutant (I and J) or the wild-type sequence (K and L). At their 3' end primers I and K harbor bases matching the mutant (green) or wild-type sequence respectively. Primers I and K were designed with melting-temperature properties corresponding to primers J and L for optimal PCR amplification. All transformants were also verified by PCR (using primers B and D) and sequencing of the resulting amplicon with the oligonucleotide (primer M) listed in Table S2.

**B**. Genotyping of $H^{aaa}$. Genotyping of $H^{aaa}$ for the lack of residual wild-type sequences (homoplasmy of the mutation) is shown as a representative example. Primers K and L (Table S2) were used to amplify the wild-type sequence. This PCR amplification is highly sensitive as shown by the presence of an amplified fragment when a 1/100 dilution of wild-type DNA is added to a sample of $H^{aaa}$ #2 DNA.

#### Figure S3

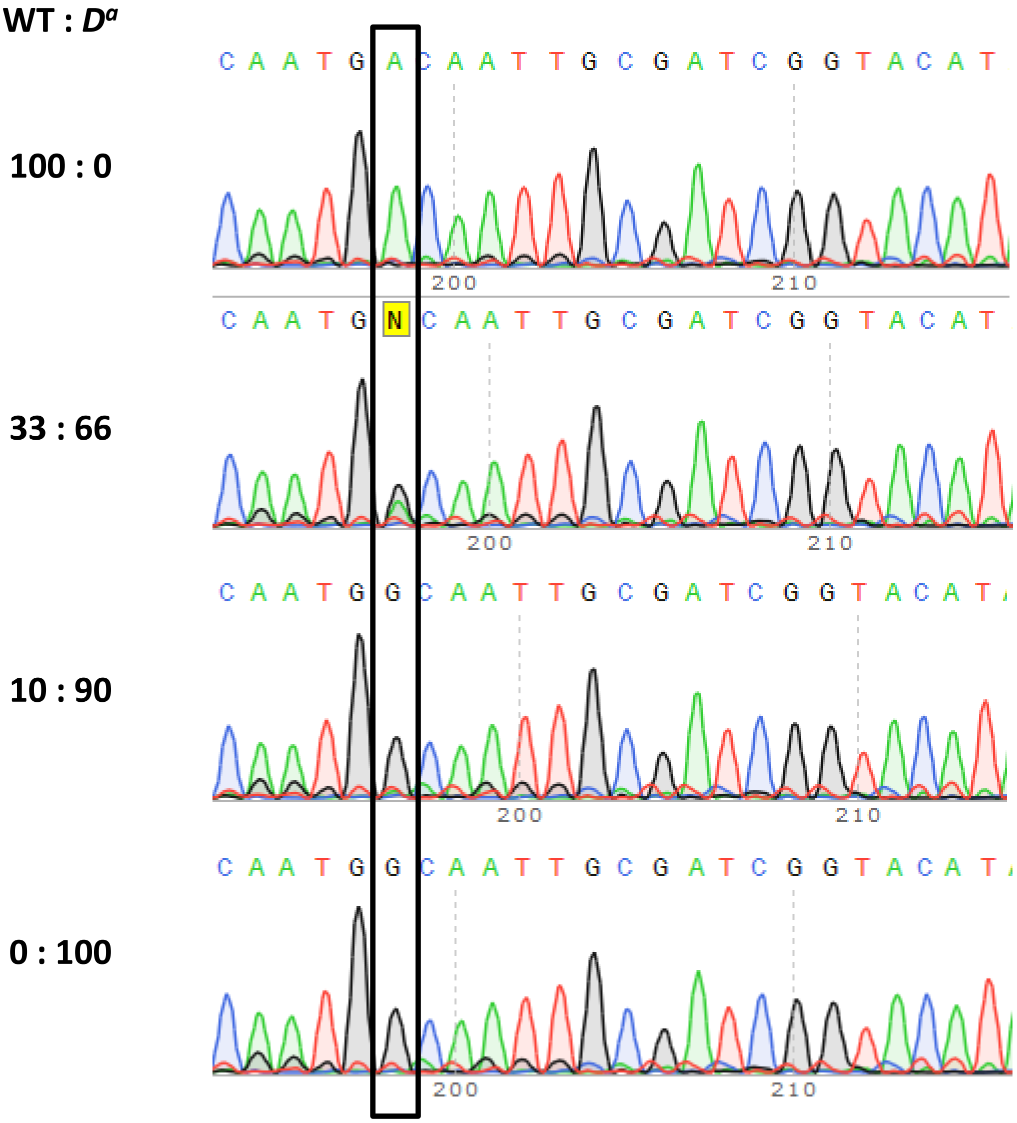


**Figure S3. Sequencing profiles of wild-type and mutant DNA mixtures.**
In cases where the PCR amplifications of the mutant versus the wild-type sequence were not specific (NS in Table S2), homoplasmy was evaluated from sequencing profiles. These strains were deemed homoplasmic when the wild-type sequence remained undetectable after at least six rounds of sub-culturing on solid media in the absence of spectinomycin selection. The limit of detection was estimated to be between 10 and 30% as follows. Mixtures of wild-type and $D^a$ genomic DNA were used as templates for amplification and sequencing (Figure S2 and Table S2). The wild-type sequence at the position boxed (A rather than G) is clearly detected when initially present in a mixture of WT : $D^a$ at 33 : 66, but not at 10 : 90.

**Figure S4**

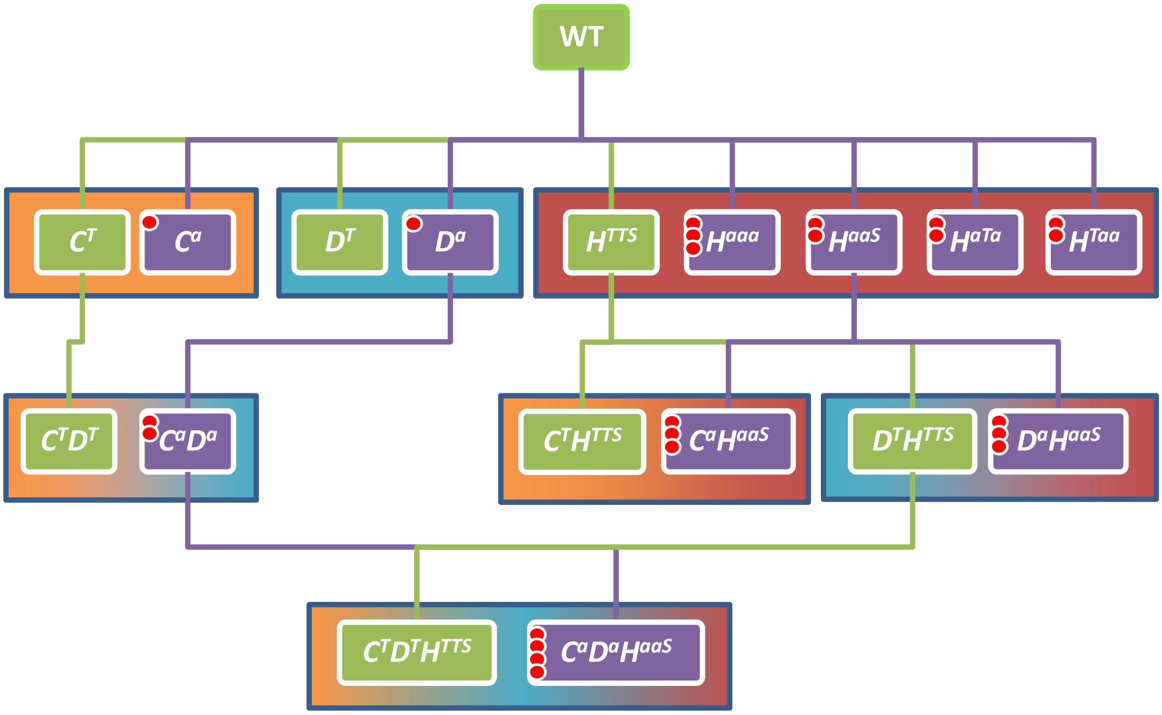


**Figure S4. Pedigree of the mutant strains.**
Strains with mutations in one, two or three chloroplast genes were obtained by repeated rounds of transformation, selection, and selectable marker removal as depicted in Fig 1. Red dots represent the total number of mutated phosphorylation sites. The control or mutant genotypes are designated by capital letters referring to the mutated protein(s) (*C*: PsbC/CP43; *D*: PsbD/D2; *H*: PsbH) with a superscript indicating whether the target phosphorylation site(s) are wild-type ($^{T}$ or $^{S}$), or have been mutated to alanine ($^{a}$). The control strains (green background) and the mutants (purple background) all carry the 483 bp footprints of the excised *aadA* cassettes.

### Figure S5

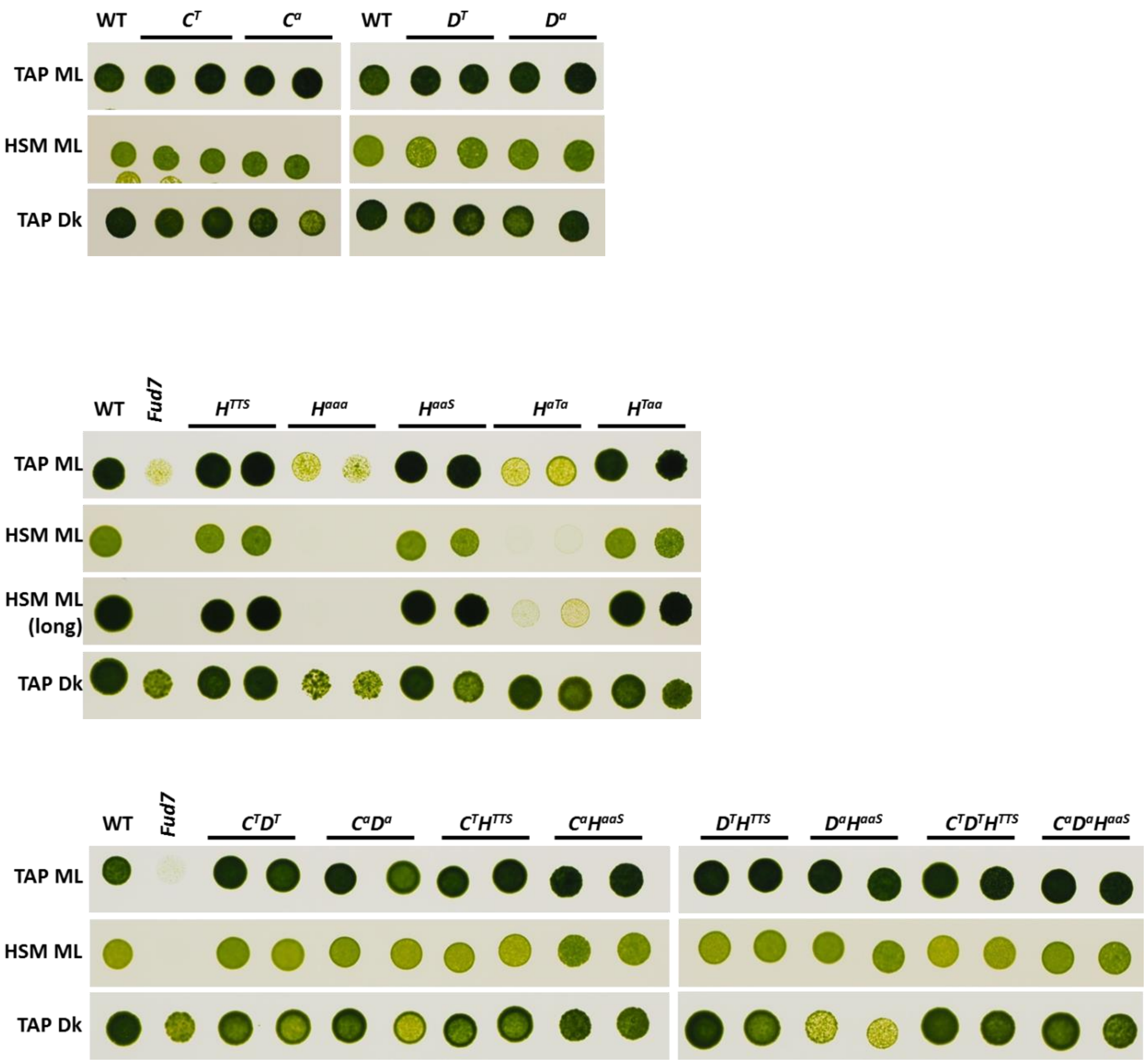


**Figure S5. Growth tests.**
Phototrophic growth was monitored on minimal medium under 60 µE $m^{-2}$ $s^{-1}$ for 5 days (HSM ML) or 19 days (HSM ML long), mixotrophic growth on acetate medium under 60 µE $m^{-2}$ $s^{-1}$ for 3 days (TAP ML) and heterotrophic growth on acetate medium in the dark for 10 days (TAP Dk). Two independent lines are shown for each genotype. A mutant with a deletion of *psbA* (*Fud7*) is shown as a non-photosynthetic control.

**Figure S6**

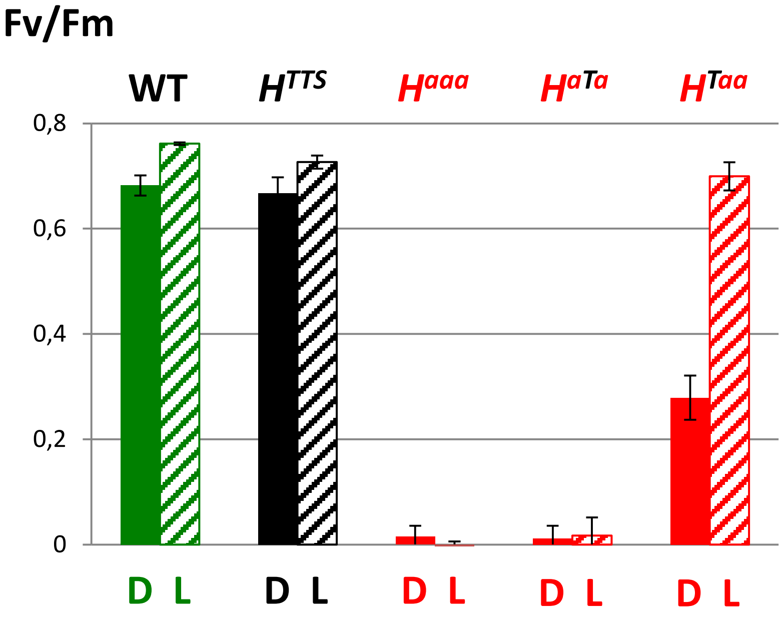


**Figure S6. Maximum quantum yield of PSII in PsbH mutants after growth in the light.**
Maximum quantum yield of photosystem II (Fv/Fm = (Fm-Fo)/Fm) after growth in the light (L) compared to growth in the dark (D; values reproduced from Fig 2A). Cells were grown in TAP medium, each value is the average of three measurements on two independent lines of identical genotype , +/- SD.

#### Figure S7

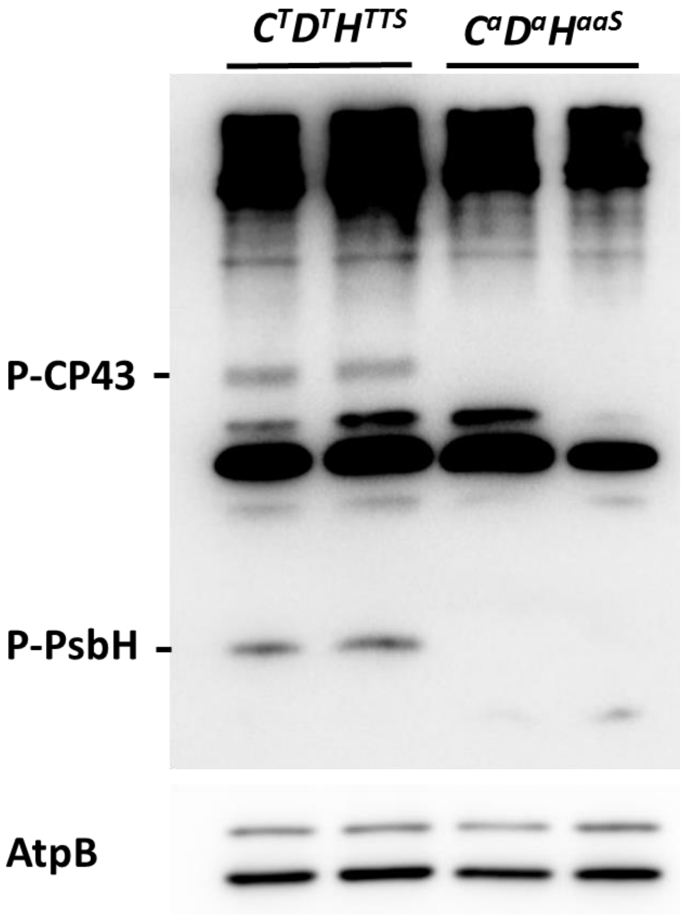


**Figure S7. Immunoblot analysis of protein phosphorylation.**
Thylakoid membrane proteins from cells treated for 1h with medium light (60 µE $m^{-2}$ $s^{-1}$) were separated by neutral LDS-PAGE followed by immunoblotting with antisera against phospho-threonine or AtpB as a loading control.

### Figure S8

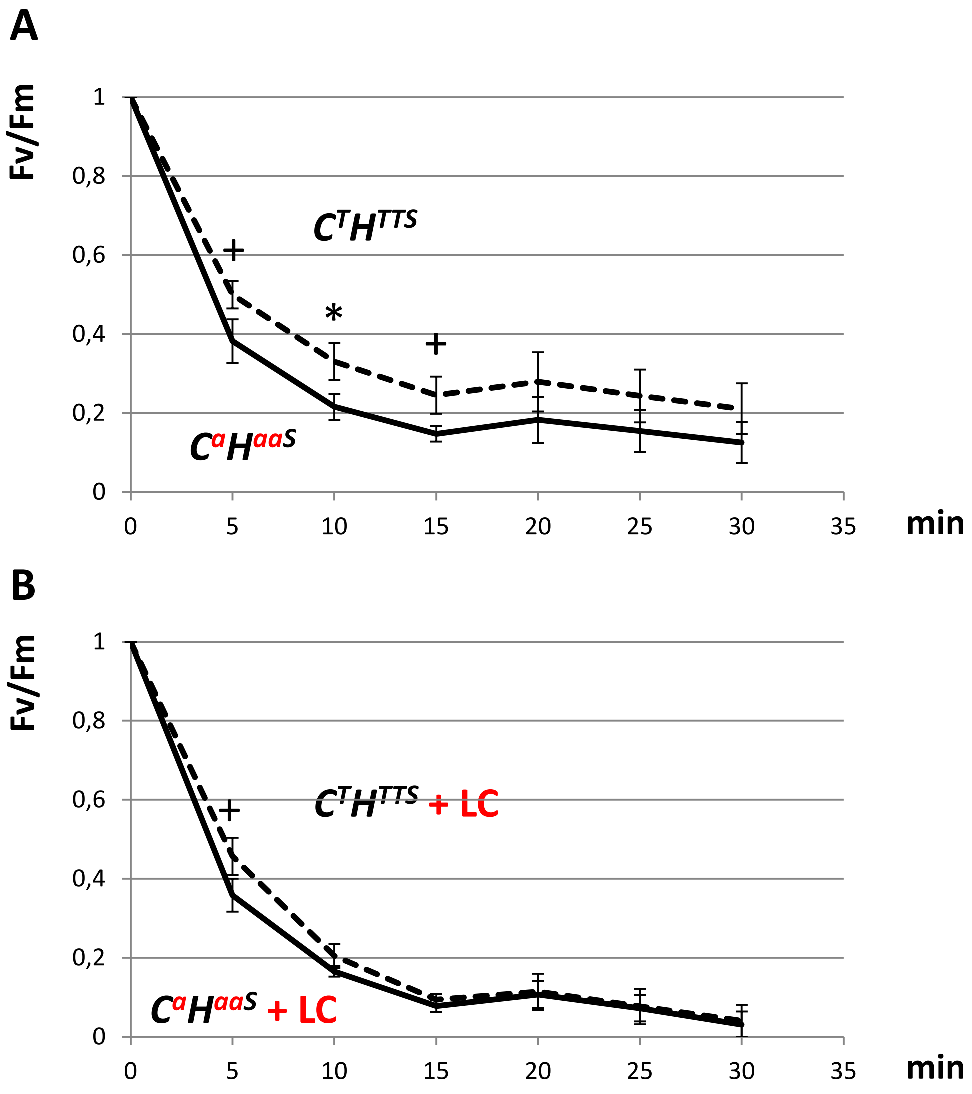


**Figure S8. Effect of translation inhibitors on photoinhibition.**

Time-course of photoinhibition at 1450 µE $m^{-2}$ $s^{-1}$ of $C^{a}H^{aaS}$ compared to the $C^{T}H^{TTS}$ control. The maximum quantum yield of PSII ($F_v/F_m$) is normalized to its value before the treatment (t = 0). Each time-point is the average of three measurements on two independent lines of identical genotype (+/- SD). Significant differences at each time point were determined using repeated measures ANOVA and a post-hoc test with Bonferroni's correction; +: p<0.008; *: p<0.0016).

A. Timecourse in the absence of inhibitors. $C^{T}H^{TTS}$ was significantly different from $C^{a}H^{Taa}$ in the absence of inhibitors (repeated measures ANOVA between the genotypes, p=0.00535)

B. Timecourse in the presence of inhibitors of chloroplast translation (+LC; 500 µgm$L^{-1}$ lincomycin and 100 µgm$L^{-1}$ chloramphenicol). $C^{T}H^{TTS}$ was not significantly different from $C^{a}H^{Taa}$ .
